# Three annotated tiger beetle genomes (Coleoptera, Adephaga, Cicindelidae)

**DOI:** 10.64898/2026.07.16.739005

**Authors:** Jamie Ramirez, Ming-Hsun Chou, Grey T. Gustafson

## Abstract

Advances and accessibility to next-generation technology provide opportunities to sequence non-model organisms. Despite this increase in whole-genome sequencing data, annotation and the production of reference genomes remain limited. Reference genomes are a critical tool for a variety of studies in evolutionary biology, functional genomics, and conservation genetics. Tiger beetles (Cicindelidae) are a diverse and globally distributed family of beetles that serve as bioindicator taxa and flagship species for insect conservation. Here, we report highly complete, contiguous, and annotated genome assemblies representing draft reference genomes for three species of tiger beetle spanning the phylogeny. These draft reference genomes are for Audouin’s night-stalking tiger beetle, *Omus audouini*; the montane giant tiger beetle, *Amblycheila baroni*; and the western red-bellied tiger beetle, *Cicindelidia sedecimpunctata*.

**Article Summary:** Tiger beetles are a charismatic group with ∼3000 species distributed globally. Despite their popularity among insect enthusiasts and their role as bioindicators of ecosystem health, the group currently lacks a reference genome. This article outlines genome assembly and annotation for three tiger beetle species that span evolutionary relationships within the lineage. Quality control analyses show that the assemblies are reference quality and demonstrate high contiguity, completeness, and accuracy. The resulting draft genome annotations will be a valuable resource for scientific endeavors and allow for continued research on tiger beetles and their allies.

## Introduction

Recent advances in high-throughput sequencing technology, especially long-read sequencing, have resulted in greater accuracy, power, and affordability than ever before (van Dijk et al. 2023; Satam et al. 2023; Wetterstrand 2023; Mahmoud et al. 2025; Ruckman and Long 2026). Notable initiatives like the Earth BioGenome Project, the i5k Initiative, and regional projects like the Darwin Tree of Life have capitalized on this aspect, culminating in a substantial increase in available genomic sequence data. Among the data being generated are those for non-model organisms, particularly with regard to reference genomes (Formenti et al. 2022). Reference genomes are assemblies that are highly complete, contiguous, and annotated (Formenti et al. 2022), allowing them to function analogously to type specimens, serving as a standard genome for a species (Ballouz et al. 2019). Reference genomes are an essential resource in biological research, providing a tool for reference-based assembly, variant calling, sequence alignment, gene annotation, and functional analyses, among others (Formenti et al. 2022; Ruckman and Long 2026). With regard to non-model organisms, reference genomes are a critical tool for conservation efforts, where variant calling allows estimation of heterozygosity and accurate quantification of inbreeding depression from a genome-wide perspective (Kardos et al. 2016; Formenti et al. 2022). This is in addition to the ability to assess gene introgression and hybridization (Twyford and Ennos 2012). However, annotation, a difficult yet critical aspect for producing reference genomes, remains a limiting factor forming a bottleneck in this time of advanced whole-genome sequence generation (Mora-Márquez et al. 2021; Ruckman and Long 2026).

Tiger beetles (Cicindelidae) are a diverse family of terrestrial, predatory beetles with nearly 3,000 species (Wiesner 2020). They have a global distribution, being absent only from arctic regions and highly remote islands, with their highest diversity in the tropics (Cassola and Pearson 2000). Like the closely related ground beetles (Carabidae), tiger beetles can serve as bioindicator taxa for ecosystem health (Pearson and Cassola 1992; Costa and Zalmon 2019), and their charisma makes them an excellent flagship species for insect conservation (Cassola and Pearson 2000; Pearson and Cassola 2007). Furthermore, many tiger beetle species themselves are of conservation concern (Fernández et al. 2026), with nearly a third of the recognized taxa in the United States of America identified as being threatened, and some already extinct (Knisley et al. 2014). Despite their popularity, studies on their evolution using molecular data (reviewed in Gough et al. 2019) have lagged behind related groups like the predatory aquatic beetles (Short 2018; Baca et al. 2025; Bergsten et al. 2025). Just recently (May 2026), the first whole-genome sequence data were published in the National Center for Biotechnology Information (NCBI) Sequence Read Archive (SRA) for any tiger beetle (BioProject PRJNA1348490). These data are primarily low-coverage, short-read (‘genome skimming’) Illumina NovaSeq data generated as part of the first phylogenomic inference of the family with complete tribal-level sampling (Gustafson et al. 2026). Thus, Cicindelidae is presently without highly complete, contiguous, and annotated genomes that can serve as reference genomes. Here, we generate three draft reference genomes for taxa distributed across the tiger beetle phylogeny (Fig. 1) to aid future studies on the evolution and conservation of this notable group.

**Figure 1.**
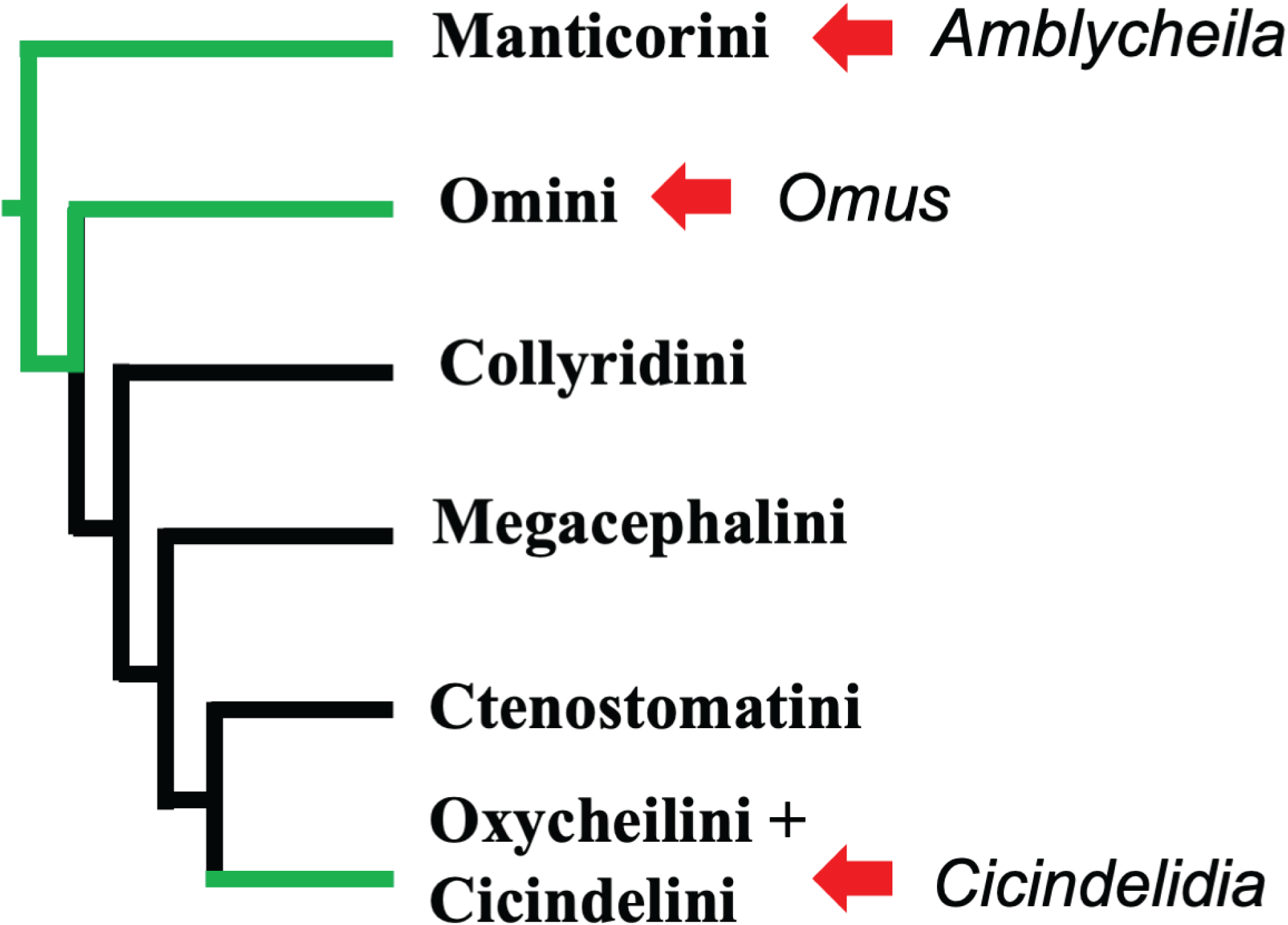
Phylogeny for the family Cicindelidae based on Gustafson et al. (2026). Red arrows denote placement of *Omus*, *Amblycheila*, and *Cicindelidia* within the lineage. Green lines denote portions of the lineage represented by reference genomes.

## Materials and methods

### Sequencing

Individuals of: *Omus audouini* (NES728), Audouin’s night-stalking tiger beetle; *Amblycheila baroni* (NES739), the montane giant tiger beetle; and *Cicindelidia sedecimpunctata* (NES740), the western red-bellied tiger beetle, were collected alive from the field (S1 Table 1) and maintained in the laboratory before transport for sequencing. The night before shipping, individuals were placed in a −80 ℃ freezer for dispatchment, then shipped the next day with dry ice. Specimens were first received by lab technicians at Novogene Co. Inc. in Irvine, California, USA, for DNA extraction, quantification, and Pacific BioSciences (hereafter PacBio) DNA high-fidelity (HiFi) library preparation, then sent to the Beijing Novogene branch for third-generation PacBio HiFi sequencing on a full lane of the PacBio Revio system.

**Table 1.** Summary table of genome assembly statistics including total length, contig number, BUSCO completeness, contiguity, accuracy, and number of predicted genes for NES728, NES739, and NES740.

| NES | Taxon | Total Length (Mbp) | Contig Number | After Decontamination BUSCO % C | N50 (Mbp) | L50 |
| --- | --- | --- | --- | --- | --- | --- |
| 728 w/ Chordata | *Omus audouini* | 938 | 533 | 97.2 | 6.76 | 32 |
| 728 w/out Chordata | *Omus audouini* | 881 | 519 | 92.4 | 6.33 | 32 |
| 739 | *Amblycheila baroni* | 1086 | 255 | 97.48 | 15.19 | 22 |
| 740 | *Cicindelidia sedecimpunctata* | 819 | 1780 | 96.7 | 33.05 | 9 |
| **Avg** |  | 928.67 | 851.33 | 95.53 | 18.19 | 21 |

| NES | N90 (Kbp) | Unmapped Positions % | Error per Mb % | High read support % (>81) | OrthoDB AA Annotation Completeness % | OrthoDB Predicted Genes |
| --- | --- | --- | --- | --- | --- | --- |
| 728 w/ Chordata | 795.9 | 4.19E-05 | 7.23E-06 | 0.67 | 92.7 | 39009 |
| 728 w/out Chordata | 778.9 | 4.56E-05 | 8.61E-06 | 0.68 | 88.4 | 35114 |
| 739 | 3287.4 | 8.01E-05 | 5.22E-06 | 0.81 | 96 | 27272 |
| 740 | 92 | 5.66E-06 | 2.28E-04 | 0.88 | 93.7 | 27564 |
| **Avg** | 1386.1 | 4.38E-05 | 8.06E-05 | 0.79 | 92.70 | 29983.33 |

### Read Statistics and Coverage Estimation

Genome size estimation and heterozygosity analyses were performed prior to assembly using a histogram generated with Jellyfish v 2.3.1-6 (Marcais and Kingsford 2011) with a k-mer length of 21 nucleotides, as input to Genome Scope 2.0 with default settings for diploid organisms (T. Rhyker Ranallo-Benavidez et al. 2020). Results were visualized using the Genome Scope online platform (http://genomescope.org/genomescope2.0/). Read length distribution plots and N50 statistics were generated on HiFi reads using assembly-stats v 1.0.1-10 (Trizna 2020), bioawk (Shen et al. 2024), ggplot (Ginestet 2011), and custom scripts (Kim and Kim 2022). Genome coverage was estimated as the difference between total base pairs and haploid genome length for each genome (S1 Table 1).

### Genome Assembly and Quality Control

HiFi reads were filtered with HiFiAdapterFilt v 3.0.1 (https://github.com/sheinasim-USDA/HiFiAdapterFilt/blob/master/README.md) and assembled with HiFiasm v 0.25 and v 0.21 using default settings. Contig coverage length distributions for each assembly, which provide a visual representation of N50, were generated using scripts from Kim and Kim (2022).

To identify heterozygosity and remove ‘junk’ haplotigs, purge_haplotigs (Roach et al. 2018) was employed to construct a histogram and identify the need for purging. If the histogram displayed two peaks, one for diploid and a second for haploid, then purging was performed before decontamination. Purging cutoff values were determined following a visual inspection of the coverage histograms as outlined in the “Manual Step” for the Purge Haplotig pipeline. Cutoff values are provided to analyse individual contigs and produce a CSV file containing potential flagged haplotigs for removal using BEDtools or reassignment with minimap2 during the final purging step (Roach et al. 2018). This process was performed in two separate analyses on sample NES740 to investigate and confirm the duplication levels observed in downstream analyses. Both assemblies in which only one diploid peak was observed (purging did not occur) and the resulting purged assemblies were provided to the BlobTools2 (Challis et al. 2020) software for decontamination (outlined below). BUSCO v 6.0 (Tegenfeldt et al. 2025) analyses were generated to assess completeness (Table S1) using the coleoptera_odb12 database.

Assemblies were evaluated for potential contamination by performing BLAST+ v 2.13 using Blastn (Camacho et al. 2009) and the NCBI core_nt database. BLAST+ v 2.13 results were analyzed in BlobTools2 to identify non-arthropod or unidentifiable (i.e., ‘no-hit’) sequence contamination for removal. The assembly fasta was indexed and converted into a sorted.bam file using SamTools v 1.15.1 (Li et al. 2009), and MiniMap2 (Li 2018). The resulting BLAST+ v 2.13 query output file, sorted.bam file and BUSCO results for the associated assemblies were provided to BlobTools2 for database construction and BlobPlot generation to visualize contamination levels. Summary text outputs from BlobTools were provided to Seqkit v 2.13 (Shen et al. 2024) to perform decontamination and generate a polished assembly. Following targeted filtering, a second BlobPlot was generated to validate performance and determine whether additional filtering was necessary. BUSCO analysis using the Coleoptera_odb12 database was conducted again to compare with earlier results.

Summary text outputs were assembled using BlobTools2 and Seqkit v 2.13. When necessary, these outputs were used to construct a custom database with putative contaminated contigs for additional BLAST taxonomic profiling using the NCBI core_nt database supplemented with *Omus dejeanii* (NES11) and *Cincindela sexguttata* (GTG001) whole-genome sequence data available from NCBI BioProject PRJNA1348490 (referred to as the “updated database”; Camacho 2008). Quast (Gurevich et al. 2013) analyses were conducted on the decontaminated assemblies to complement BUSCO results and provide additional information on assembly contiguity. Accuracy of final assemblies was assessed using the program Referee (Thomas and Hahn 2019).

### Repeat Region Identification and Masking

Assemblies were soft masked for known metazoan repeats using RepeatMasker 4.2.2 (Smit et al. 2013). Species-specific repeats within each genome were identified, and a de novo repeat library was constructed using RepeatModeler 2.0.7 (Flynn et al. 2020). The identified novel repeats and subsequent de novo libraries were used to soft-mask the initial metazoan-masked assemblies (referred to as the SMM assembly).

### Annotation

Two separate genome annotations were performed using two reference sources. The first approach used proteins of closely related ground beetles: *Carabus problematicus*, *Nebria brevicollis*, and *Pterostichus madidus*, from the Darwin Tree of Life project hosted by Ensemble (Dyer et al. 2025) (referred to as “curated carabid protein hints”). The second approach used the arthropod proteins from the OrthoDB v12 protein database (Kuznetsov et al. 2023). Protein-coding gene prediction was performed using GeneMark-ETP v 1.0 (Brůna et al. 2024), and AUGUSTUS v 3.5.0-9 (Stanke et al. 2004) within the BRAKER3 (Brůna et al. 2021; v 3.0.8) workflow on the SMM assemblies. Output protein predictions were generated for each of the associated reference sources. Completeness analyses of the resulting gene predictions from each BRAKER3 run were assessed using BUSCO v6 with the coleoptera_odb12 database and protein mode settings.

## Results and Discussion

### Sequencing, Read Statistics, and Coverage Estimation

Preliminary read analysis (S1 Table 1; S2a–c ) showed that sequencing yielded 2.1, 2.95, and 2.97 million reads. Read statistics recovered an average N50 of 17006.3 bp and N90 of 12716 bp across all genomes. Statistical analysis of the PacBio HiFi sequence data recovered yields of 33.98 Gbp (NES728), 45.7 Gbp (NES739), and 44.3 Gbp (NES740), with an average of 41.3 Gbp (Table S1; S2). The average number of reads across the three datasets is 2.67million, with NES740 possessing the most (2974806), and NES728 the least (2095826) number of reads.

GenomeScope analyses show high levels of homozygosity (97.25-100%) and low levels of heterozygosity (0-2.75%) in the reads (S1 Table 1; S3–5). The lowest level of heterozygosity was found in NES739 (S1 Table 1; S4e), Amblycheila baroni, at 0.8%. NES728 (S1 Table 1; S3e) had 0-2.16% and NES740 (S1 Table 1; S4e) 2.72-2.75% heterozygosity. Estimated coverage is ∼32x, ∼69.6x, and ∼76.7x for NES728 (S3), NES739 (S4), and NES740 (S5), respectively.

### Genome Assembly and Quality Control

Following assembly, initial BUSCO analyses using the Coleoptera_odb12 lineage dataset obtained high completeness scores of 97.5%(NES728), 97.5% (NES739), and 96.8% (NES740), comprising 3635, 3635, and 3608 complete BUSCOs out of the 3729 BUSCOs searched (Table S1).

Histograms generated using purge_haplotigs (S6a–c) recovered one peak (diploid) for NES728 (S6a) and two peaks for NES739 (S6b) and NES740 (S6c; S7a–c). The latter two samples proceeded through the purge_haplotigs pipeline as described above. This process was performed in two separate analyses on sample NES740 ( S6c; S7a–c ) to investigate and confirm the duplication levels observed in downstream analyses, which yielded comparable results (S7b–c). After purging, BUSCO results (referred to as “Before Decontamination” or BD) showed completeness scores consistent with those from the initial BUSCO analyses provided above (S1 Table 1).

BlobTools blob plots upstream of the decontamination stage revealed conflicting GC content and spikes in coverage characteristic of contamination and differing from the main genome (S8–9). NES739 had the lowest amount of contamination from one source (no hit; S9a), while NES740 was afflicted with low levels of contamination from five sources (S9c). Conversely, NES728 experienced high levels of contamination, predominantly from bacteria, fungi, and unidentified sources (S8a, S8c). This result could be attributed to differences in the timing of specimen preparation for shipping or to residual contamination from field collection.

Following decontamination (referred to as “After Decontamination” or AD), Blob Plots and BUSCO scores were compared with earlier BD results (Fig. 2; Table 1; S1 Table 1; S8a–d; S9a–d; S10a–d; S11a–d). Blob plots demonstrated the removal of all specified contamination phyla and the absence of the need for additional quality control measures. However, when Chordata was filtered for NES728 using BlobTools, BUSCO scores dropped by 5.12% (S1 Table 1; S10a–d). This difference may stem from the underrepresentation of tiger beetles in the NCBI database and difficulty identifying orthologs for non-model organisms, or difficulty in identifying incomplete BUSCO contigs following decontamination using BlobTools. In response to the differing BUSCO completeness scores, the 14 contigs for NES728 w/ Chordata were used as input (S12) for a secondary BLAST inquiry against the updated database (S13). The resulting summary text file identified the 14 putative Chordata contigs as Arthropoda and with the local tiger beetle sequences (S13, S14). Altogether, these results suggest the “Chordata contamination” was misidentified due to a lack of tiger beetle reference sequence data. We retained this assembly (w/ Chordata) alongside the w/out Chordata assembly for further analysis.

**Figure 2.**
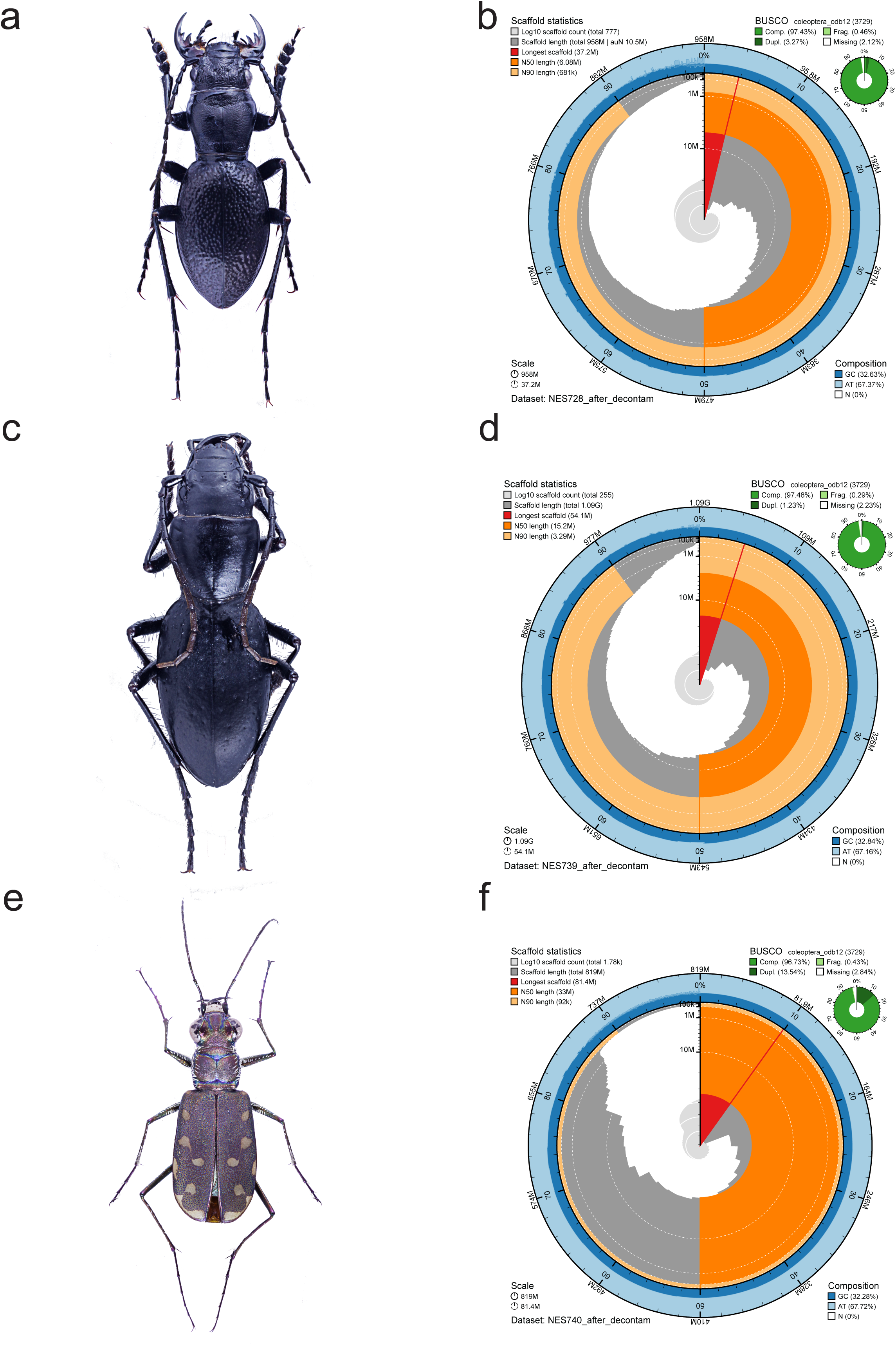
Snailplots generated by BlobTools show BUSCO completeness and contiguity (N50 and N90) results for *Omus audouini* (NES728, Fig.2 a–b), Audouin’s night-stalking tiger beetle; *Amblycheila baroni* (NES739, Fig. 2c–d), the montane giant tiger beetle; and *Cicindelidia sedecimpunctata* (NES740; Fig. 2e–f), the western red-bellied tiger beetle

Review of AD blob plots demonstrated the successful removal of impurities(S8a–d; S9a–d). As a whole, differences in BD and AD BUSCO completeness scores were largely negligible for NES728 w/ Chordata, NES739 and NES740 (-/+ 0.01%; S1 Table 1; S10b, S11b, S11d).

The BUSCO AD (Table 1; S1 Table 1; S10–11) evaluation yielded an average completeness score of 95.9% across all genomes, demonstrating high genome assembly quality. The highest completeness scores were observed for NES739 (97.48%; Table1; S1 Table 1; S11b), the second highest seen in NES728 w/ Chordata (97.2%; S10b), followed by NES740 (96.73%; S11d), with the lowest exhibited by NES728 w/out Chordata (92.4%; Table 1; S1 Table 1; S10d). Low levels of missingness are reported for all samples, with an average of about 3.58%, the lowest seen in NES739 (S1 Table 1; S11b), and the highest observed in NES728 w/out Chordata (7.11%, S1 Table 1; S10d). Scaffold N50 values vary across all samples, but all suggest high contiguity (Table 1; S1 Table 1). The lowest N50 value is observed in NES728 w/ Chordata (6.08Mb; Table 1; S1 Table 1; S10b), second lowest is NES728 w/out Chordata (6.33Mb; Table 1; S1 Table 1; S10d), the second highest by NES739 (15.2Mb; Table 1; S1 Table 1; S11b), and the highest N50 value belongs to NES740 (33Mb; Table 1; S1 Table 1; S11d). Additionally, high N90 values were exhibited in NES728 w/out Chordata (779Kb; Table 1; S1 Table 1; S10d), NES728 w/ Chordata (681Kb); Table 1; S1 Table 1; S10b) and NES739 (3.29Mb; Table 1; S1 Table 1; S11b). In contrast, the lowest N90 value was seen for NES740 (92Kb; Table 1; S1 Table 1; S11d), potentially due to scaffold fragmentation or genome repeats influencing assembler performance.

Quast analyses (S1 Table 1; S15, S16) recover genome assemblies near 1Gb, with lengths of 881 Mb (NES728 w/out Chordata; Table 1; S1 Table 1; S15a, S15c, S15e, S15g, S15i), 938 Mb (NES728 w/ Chordata; Table 1; S1 Table 1; S15b, S15d, S15f, S15h, S15j), 1.1 Gb (NES739; Table 1; S1 Table 1; S16a, S16c, S16e, S16g, S16i), and 819 Mb (NES740; Table 1; S1 Table 1; S16b, S16d, S16f, S16h, S16j) reported. These assemblies are represented by 519 (NES728 w/out Chordata; Table 1; S1 Table 1; S15a), 533 (NES728 w/ Chordata; Table 1; S1 Table 1; S15b), 255 (NES739; Table 1; S1 Table 1; S16a), and 1780 (NES740; Table 1; S1 Table 1; S16b) contigs. N50 values are consistent in both BUSCO and Quast analyses, suggesting highly contiguous assemblies (S1 Table 1). The highest N50 and lowest L50 values were reported for NES740 (N50:33.05 Mbp; L50:9; S1 Table 1; S16b, S16d). In contrast, the lowest N50 and highest L50 were found to belong to NES728 (w/out Chordata: N50: 6.33 Mbp; L50: 32; Table 1; S1 Table 1; S15a; S15c; w/ Chordata: N50: 6.76 Mbp; L50: 32; Table 1; S1 Table 1; S15b; S15d), with intermediate values observed in NES739 (N50: 15.19 Mbp; L50 22; Table 1; S1 Table 1; S16a; S16c). The highest N90 and lowest L90 values are seen in NES739 (∼3.29 Mbp; L90: 79; Table 1; S1 Table 1; S16a; S16c), the lowest N90 and highest L90 in NES740 (N90: 92 Kbp; L260; Table 1; S1 Table 1; S16b; S16d), and intermediate results belonging to NES728 (w/out Chordata:N90: 778.9 Kbp; L90: 209; Table 1; S1 Table 1; S15a; S15c; w/ Chordata: N90: 795.9 Kbp; L90: 214; Table 1; S1 Table 1; S15b; S15d). Area under the Nx curve (auN) supplements N50 and N90 by providing a measure of assembly contiguity that weighs larger contigs more heavily than shorter contigs (Gurevich et al. 2013; Li 2020; Quast manual: https://quast.sourceforge.net/docs/manual.html). The auN values surpass the N50 values for all assemblies, further supporting high-quality assemblies (S1 Table 1; S15a–f; S16a–f).

Referee analyses assessed the accuracy of the assemblies by identifying erroneous or supported sites and providing confidence scores (Thomas and Hahn 2019). Total length and number of contigs were consistent across previous completeness and contiguity analyses for all assemblies. Unmapped positions were very low for all assemblies, with the highest value observed in NES739 (8.01E-05; Table 1; S1 Table 1), intermediate values in NES728 w/out Chordata (4.56E-05; Table 1; S1 Table 1) and NES740 (5.66E-06; Table 1; S1 Table 1), and the lowest value in NES728 w/ Chordata (4.19E-05; Table 1; S1 Table 1). The highest errors corrected per million base pairs are observed in NES740 (228; Table 1; S1 Table 1), the lowest in NES739 (5.22; Table 1; S1 Table 1), and the moderate values belong to NES728 w/out Chordata (8.6; Table 1; S1 Table 1) and NES728 w/ Chordata (7.23; Table 1; S1 Table 1). The high duplication rate observed in NES740 may be affecting base-call accuracy, contributing to the higher error rates in this sample. The assembly with the highest proportion of confidence scores (Bins 81–91) was observed for NES740 (0.88; Table 1; S1 Table 1), and the lowest was seen in NES728 w/ and w/out Chordata (0.67–0.68; Table 1; S1 Table 1), with NES739 (0.81; Table 1; S1 Table 1) possessing the second highest value. Median values (Bins 51-80) were highest for both NES728 assemblies (0.19; Table 1; S1 Table 1), second highest in NES739 (0.12; Table 1; S1 Table 1), and lowest in NES740 (0.04; Table 1; S1 Table 1).

To summarize, statistical analyses of the AD draft genomes, including BUSCO completeness scores, N50, L50, N90, L90, auN, and Referee results, suggest highly complete, contiguous, and accurate assemblies. High contiguity is further demonstrated with all three genomes being predominantly composed of a small number of large contigs (N50 and auN values; Table 1; S1 Table 1; S9–10). Further comparison of the N50, L50, auN, N90, and L90 values for NES728 and NES740 shows a decline in contig size and an increase in contig quantity. These results indicate a significant contribution from shorter contigs following the N50 threshold for the aforementioned samples. However, this does not compromise the integrity of the assembly quality observed across these aforementioned assemblies, as Referee results demonstrate a low percentage of unmapped positions. Additionally, Referee results suggest that NES740, though exhibiting higher duplication levels, which may have contributed to the highest error values observed, also has high read support across most of the assembly. The NES728 Referee results show that the assembly has relatively low correction and error rates, with confidence scores spanning median to high values. Finally, completeness and contiguity scores for NES739 suggest balanced contig distribution, with larger contig sizes observed in both N50 and N90 values. NES739 also demonstrates low unmapped positions, error rates, and predominantly high read confidence values.

### Annotation

Annotation using the OrthoDB12 conserved protein hints performed better for NES728 (w/ Chordata: 39009 genes; w/out Chordata: 35114; Table 1; S1 Table 1) than the curated carabid protein hints (w/ Chordata: 37499 genes; w/out Chordata: 34567; Table 1; S1 Table 1). In contrast, both NES739 and NES740 yielded more predicted genes when using the curated carabid protein hints (27766 and 29100 genes, respectively) than with the OrthoDB12 hints (27272 and 27564 genes; Table 1; S1 Table 1). However, both approaches are systematically limited due to the lack of representation of closely related cicindelid taxa.

BUSCO analysis of both annotated gene prediction datasets using the curated carabid and OrthoDB12 protein hints yielded comparable completeness evaluation results, with the OrthoDB12 hints performing slightly better (Table 1; S1 Table 1). Results from the annotated protein predictions yielded 93% (NES728), 95.8% (NES739), and 93.3% (NES740) complete genes. This includes 3003 (NES728 w/ Chordata), 2872 (NES728 w/out Chordata), 3156 (NES739), and 2709 (NES740) complete and single-copy genes, as well as 466, 436, 417, and 772 complete and duplicated genes, respectively (Table 1; S1 Table 1).

## Conclusion

The resulting assemblies have demonstrated high completeness, with an average BUSCO score of 95.5%. High contiguity was established by high N50 (average 15.33 Mbp), N90 (average 1238.6 Kbp), and auN (average 18.73 Mbp) values from both BUSCO and Quast analyses. High assembly accuracy was verified by Referee analyses, which reported low proportions of unmapped regions and error rates, along with high assembly quality scores across the majority of the genomes. Our results indicate draft genome assemblies of reference quality which will serve as a valuable tool for future scientific endeavors by Cicindelid workers.

## Funding

Funding has been provided by the National Science Foundation to G.T.G. (NSF DEB #2208620 “Manticores and Giant Tigers: Utility of UCEs for inferring shallow and deep evolutionary history within the tiger beetle tribe Manticorini (Coleoptera: Cicindelidae)”).

## Conflicts of Interest

None declared.

## Data Availability

The genome assemblies and annotations are available in the NCBI under the BioProject PRJNA1494713 and at Zenodo (https://doi.org/10.5281/zenodo.21363288).

## Supporting information

Combined_Supplementary_S1Table1_Figures

## Acknowledgements

Marc Tollis (Northern Arizona University) is thanked for advice on annotation approaches. Computational analyses were run on Northern Arizona University’s Monsoon computing cluster, funded by Arizona’s Technology and Research Initiative Fund. This research was supported by National Science Foundation grant DEB# 2208620.

---

Supplemental Table 1. Expanded table including specimen details, read statistics, Genome Scope read analyses, BUSCO scores throughout the workflow, Quast contiguity scores, Referee assembly accuracy scores, and resulting annotation statistics for NES728, NES739, NES740.

Supplemental Figure 2. Read statistics including contiguity scores and the graphs demonstrating the relationship between density and read count (y axis) and read length (x axis) for NES728 (S2a), NES739 (S2b), and NES740 (S2c).

Supplemental Figure 3. Genome Scope results for NES728 which include coverage frequency graphs (a–d), results (e) that include levels of homozygosity, heterozygosity, genome haploid length, genome repeat length, genome unique length, model fit, read error rate, and model specifics (f) for the analysis.

Supplemental Figure 4. Genome Scope results for NES739 which include coverage frequency graphs (a–d), results (e) that include levels of homozygosity, heterozygosity, genome haploid length, genome repeat length, genome unique length, model fit, read error rate, and model specifics (f) for the analysis.

Supplemental Figure 5. Genome Scope results for NES740 which include coverage frequency graphs (a–d), results (e) that include levels of homozygosity, heterozygosity, genome haploid length, genome repeat length, genome unique length, model fit, read error rate, and model specifics (f) for the analysis.

Supplemental Figure 6. Purge Haplotigs histograms for NES 728 (a), NES 739 (b), and NES740 (c).

Supplemental Figure 7. Original histogram peaks (a) for NES740 and comparison of snail plots (left b–c) which demonstrate similar duplication results despite changes in Purge Haplotig settings (right b–c).

Supplemental Figure 8. Blobplot results for NES728 (a–d) before decontamination (S8a, S8c) and after different decontamination approaches (S8b, S8d).

Supplemental Figure 9. Blobplot results for NES739 (a–b) and NES740 (c–d). Before decontamination blobplots (NES739, S9a; NES740, S9c) and after decontamination (NES739, S9b; NES740, S9d).

Supplemental Figure 10. Snail plots result of NES728 (S10a–d) for the Chordata included dataset (S10a–b), and the Chordata omitted dataset (S10c–d). Before decontamination (S10a, S10c) is compared with after decontamination (S10b, S10d).

Supplemental Figure 11. Snail plots result for the NES739 (S11a–b), and NES740 (S11c–d). Before decontamination (S11a, S11c) is compared with after decontamination (S11b, S11d).

Supplemental Figure 12. Text file of the isolated, putative Chordata contigs for NES728 w/ Chordata dataset.

Supplemental Figure 13. Blast megablast output for NES728 (w/ Chordata) using NCBI core_nt database supplemented with local tiger beetle sequences.

Supplemental Figure 14. Summary table results for NES728 (w/ Chordata) identifying putative Chordata contigs as Arthropoda.

Supplemental Figure 15. Quast results for NES728 w/ out Chordata (S15a, c, e, g, i) and w/ Chordata (S15b, d, f, h, j). Details including contiguity statistical reports (w/out Chordata S15a; w/ Chordata S15b), area under the curve graph (w/out Chordata S15c; w/ Chordata S15d), cumulative contig length graph (w/out Chordata S15e; w/ Chordata S15f), GC contig per window (w/out Chordata S15g; w/ Chordata S15h), and GC content per contigs (w/out Chordata S15i; w/ Chordata S15j) is provided.

Supplemental Figure 16. Quast results for NES739 (S16a, c, e, g, i) and NES740 (S16b, d, f, h, j). Details including contiguity statistical reports (NES739 S16a; NES740 S16b), area under the curve graph (NES739 S16c; NES740 S16d), cumulative contig length graph (NES739 S16e; NES740 S16f), GC contig per window (NES739 S16g; NES740 S16h), and GC content per contigs (NES739 S16i; NES740 S16j) is provided.

## Notes

https://doi.org/10.5281/zenodo.21363288

### Competing Interest Statement

The authors have declared no competing interest.

