## Supplementary material for "Three annotated tiger beetle genomes (Coleoptera, Adephaga, Cicindelidae)": Combined_Supplementary_S1Table1_Figures

**Specimen Details**

| NES | Taxon | Accession | Collection locality |
|---|---|---|---|
| NES728 | *Omus audouini* | JCANOZ000000000 | USA: WA: Olympia, Mima Falls Trail, 46.90677 -123.06445, 2025.V.11, Ming-Hsun Chou col. |
| NES739 | *Amblycheila baroni* | JCANOY000000000 | USA: AZ: Pima Co: Santa Catalina Mts. General Hitchcock HWY #39 32.36209, -110.71827, 5479 ft elevation, 11vii2025, NAILS11.07.2025E Col. by S. Phillip & J. Ramirez |
| NES740 | *Cicindelidia sedecimpunctata* | JCANOX000000000 | USA: AZ: Cochise County: Whetstone Mts. French Joe "Spring" 31.81025, -110.39556 ~5200 ft elevation 10vii2025 NAILS10.07.2025D Col. by S. Phillip & J. Ramirez |
| GTG001 | *Cicindelidia sexguttata* | SRR36226413 | USA: NC: Wilkes Co, Hwy 421 10 mi E Wilkesboro 1July1988 coll. Whiting & Terry |
| NES11 | *Omus dejeanii* | SAMN52902523 | OR: Multnomah Co., Mt. Tabor, 22April2020, leg. M. Kippenhan |

**Read Statistics: Before Assembly**

**N50 Read Statistics**

| Voucher # | Total bp | Number of Reads | Average | Largest | N50 | N90 |
|---|---|---|---|---|---|---|
| NES728 | 33978166653 | 2095826 | 16212.3 | 59802 | 17274, n = 786153 | 13053, n = 1692588 |
| NES739 | 45741219640 | 2952573 | 15491.99 | 59188 | 17109, n = 1086301 | 12845, n = 2300188 |
| NES740 | 44360660576 | 2974806 | 14912.12 | 50222 | 16636, n = 1077168 | 12250, n = 2291318 |
| Average | 41360015623 | 2674401.667 | 15538.80333 | 56404 | 17006.3 | 12716 |

**Genome Scope 2.0**

**Genome Scope Read analysis-Before assembly**

| NES | Homozygous % | Heterozygous % | Avg Haploid Length (bp) | Avg Repeat Length (bp) | Avg Unique Length (bp) | Estimated Coverage (Tot. read bp / avg haploid length) | |
|---|---|---|---|---|---|---|---|
| NES728 | 97.84-100 | 0-2.16 | 1058464592 | 734845077.5 | 323619514.5 | 32.10 | |
| NES739 | 99.13-99.16 | 0.836-0.873 | 656956996 | 336786942 | 320170054 | 69.63 | |
| NES740 | 97.25-97.28 | 2.72-2.75 | 578317284 | 185107864.5 | 393209419.5 | 76.71 | |
| Completeness and Quality Analyses: After Assembly | | | | | | | |
| BUSCO | | | | | | | |
| BUSCO coleoptera_odb12 Before decontamination | | | | | | | |
| NES | Completeness % | Completeness # | Complete and single-copy | Complete and Duplicated | Fragmented | Missing | N50 |
| NES728 | 97.5 | 3635 | 3510 | 125 | 17 | 77 | 5 Mbp |
| NES739 | 97.5 | 3635 | 3589 | 46 | 11 | 83 | 14 Mbp |
| NES740 | 96.7 | 3607 | 3089 | 518 | 16 | 106 | 33 Mbp |
| BUSCO coleoptera_odb12 After decontamination | | | | | | | |
| NES | Completeness % | Completeness # | Complete and single-copy | Complete and Duplicated | Fragmented | Missing | N50 |
| NES728 w/ Chordata | 97.2 | 3632 | 3511 | 121 | 18 | 79 | 6 Mbp |
| NES728 w/out Chordata | 92.4 | 3445 | 3325 | 120 | 19 | 265 | 6 Mbp |
| NES739 | 97.48 | 3635 | 3589 | 46 | 11 | 83 | 15 Mbp |
| NES740 | 96.7 | 3608 | 3102 | 506 | 16 | 105 | 33Mbp |

| Quast | | | | | | | |
|---|---|---|---|---|---|---|---|
| Quast After decontamination | | | | | | | |
| NES | Total Length (Mbp) | Total Contig No. | Coverage (Tot. Read bp/Assembly size) | GC (%) | N50 (Mbp); L50 (No. contigs) | N90 (Kbp); L90 (No. contigs) | auN Mbp) |
| NES728 w/ Chordata | 938 | 533 | 36.23 | 32.6 | 6.76; 32 | 795.9; 214 | 10.73 |
| NES728 w/out Chordata | 881 | 519 | 38.55 | 32.61 | 6.33; 32 | 778.9; 209 | 10.15 |
| NES739 | 1086 | 255 | 42.14 | 32.84 | 15.19; 22 | 3287.4; 79 | 18.29 |
| NES740 | 819 | 1780 | 54.14 | 32.28 | 33.05; 9 | 92; 260 | 35.74 |

| Referee | | | | | | | | | |
|---|---|---|---|---|---|---|---|---|---|
| Referee Quality After decontamination | | | | | | | | | |
| NES | Total Length (Positions) | Total Contig No. | Unmapped Positions | Unmapped Positions % | Errors Corrected | Errors Corrected % | Error per base (per million) | Bins 81-91 Highest Confidence | Bins 51-80 Median Confidence |
| NES728 w/ Chordata | 937857440 | 533 | 39254 | 4.19E-05 | 6782 | 7.23E-06 | 7.23 | 0.67 | 0.19 |
| NES728 w/out Chordata | 881420873 | 519 | 40203 | 4.56E-05 | 7588 | 8.61E-06 | 8.6 | 0.68 | 0.19 |
| NES739 | 1085518503 | 255 | 86963 | 8.01E-05 | 5666 | 5.22E-06 | 5.22 | 0.81 | 0.12 |
| NES740 | 819365743 | 1780 | 4635 | 5.66E-06 | 186873 | 2.28E-04 | 228 | 0.88 | 0.04 |

| Completeness and Prediction Statistics: After Annotation |
|---|
| BUSCO |

| BUSCO coleoptera_odb12 After annotation-AA protein hints | | | | | | |
|---|---|---|---|---|---|---|
| NES Carabid | Completeness % | Completeness # | Complete and single-copy | Complete and Duplicated | Fragmented | Missing |
| NES728 w/ Chordata | 93 | 3469 | 3003 | 466 | 131 | 129 |
| NES728 w/out Chordata | 88.7 | 3308 | 2872 | 436 | 115 | 306 |
| NES739 | 95.8 | 3573 | 3156 | 417 | 79 | 77 |
| NES740 | 93.3 | 3481 | 2709 | 772 | 106 | 142 |
| NES OrthoDB | Completeness % | Completeness # | Complete and single-copy | Complete and Duplicated | Fragmented | Missing |
| NES728 w/ Chordata | 92.7 | 3458 | 3099 | 359 | 137 | 134 |
| NES728 w/out Chordata | 88.4 | 3295 | 2949 | 346 | 124 | 310 |
| NES739 | 96 | 3578 | 3245 | 333 | 74 | 77 |
| NES740 | 93.7 | 3493 | 2795 | 698 | 94 | 142 |

| Predicted Gene and AA Statistics | | | | |
|---|---|---|---|---|
| Annotated Gene and Amino Acids Predictions (MB) | | | | |
| NES | Carabid GTF | Carabid AA | OrthoDB GTF | OrthoDB AA |
| NES728 w/ Chordata | 46.17 | 13.7 | 46.3 | 13.9 |
| NES728 w/out Chordata | 42.65 | 12.41 | 42.26 | 12.52 |
| NES739 | 37.4 | 10.9 | 38 | 11 |

| | | | | |
|---|---|---|---|---|
| NES740 | 39.5 | 11.8 | 39 | 11.8 |

| Predicted Genes | | |
|---|---|---|
| NES | Carabid Genes | OrthoDB Genes |
| NES728 w/ Chordata | 37499 | 39009 |
| NES728 w/out Chordata | 34567 | 35114 |
| NES739 | 27766 | 27272 |
| NES740 | 29100 | 27564 |

**a**

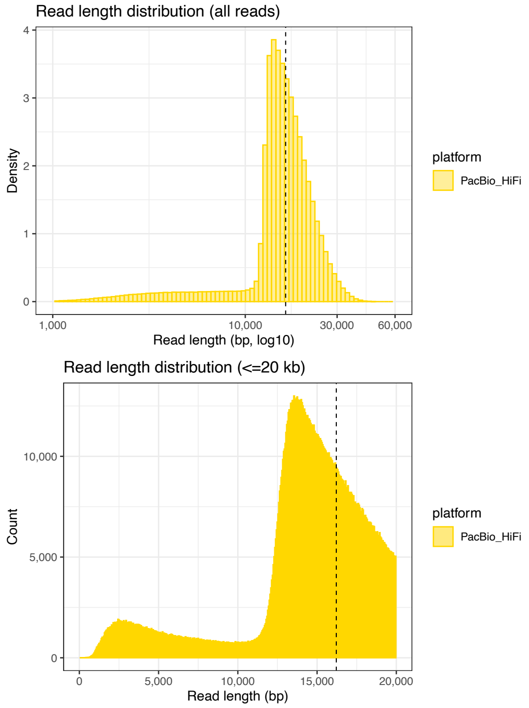

### Read Statistics

#### NES728

stats for NES728.hifi_reads.filt.fastq
sum = 33978166653, n = 2095826, ave = 16212.30,
largest = 59802
N50 = 17274, n = 786153
N60 = 16108, n = 989939
N70 = 15045, n = 1208260
N80 = 14051, n = 1441995
N90 = 13053, n = 1692588

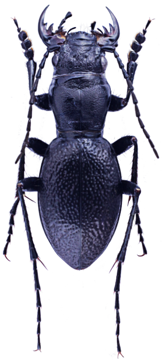

**b**

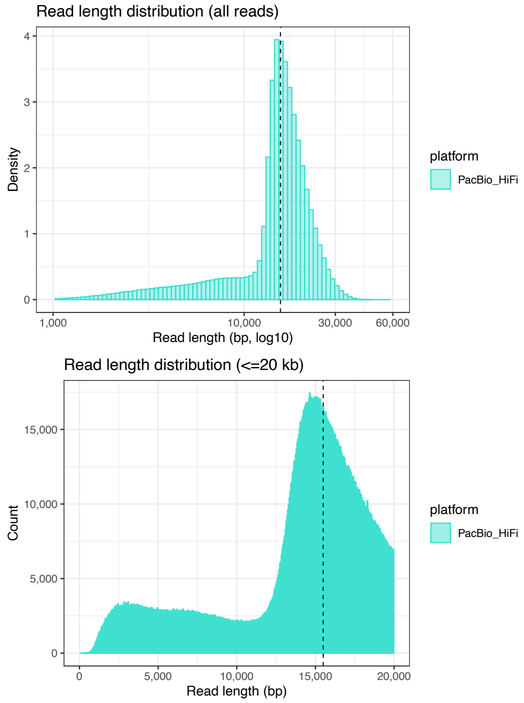

#### NES739

stats for NES739.hifi_reads.fastq
sum = 45741219640, n = 2952573, ave = 15491.99,
largest = 59188
N50 = 17109, n = 1086301
N60 = 16097, n = 1362076
N70 = 15183, n = 1654739
N80 = 14268, n = 1965311
N90 = 12845, n = 2300188

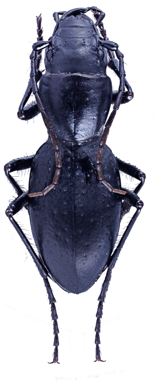

**c**

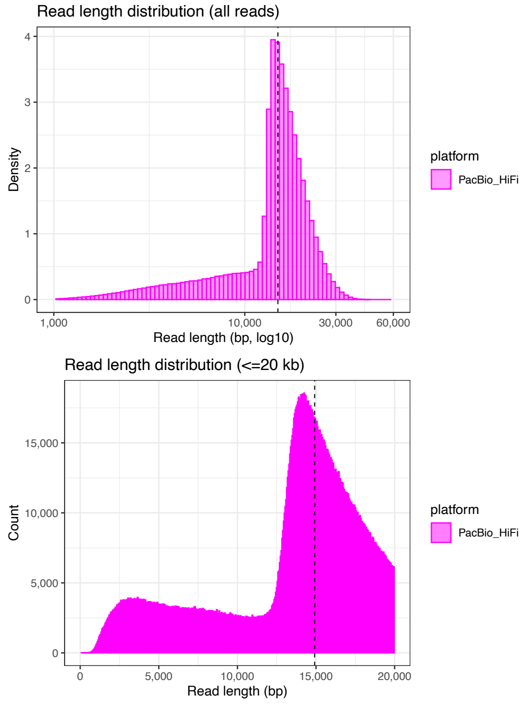

#### NES740

stats for NES740.hifi_reads.fastq
sum = 44360660576, n = 2974806, ave = 14912.12,
largest = 50222
N50 = 16636, n = 1077168
N60 = 15608, n = 1352611
N70 = 14697, n = 1645609
N80 = 13839, n = 1956584
N90 = 12250, n = 2291318

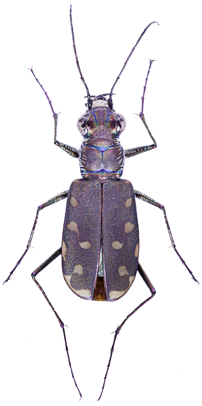

### NES728 GenomeScope

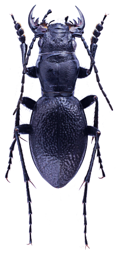

**a**

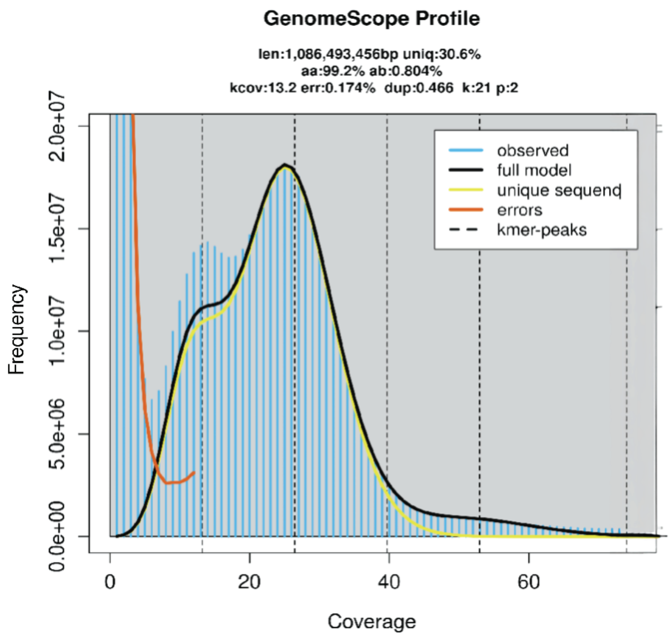

**b**

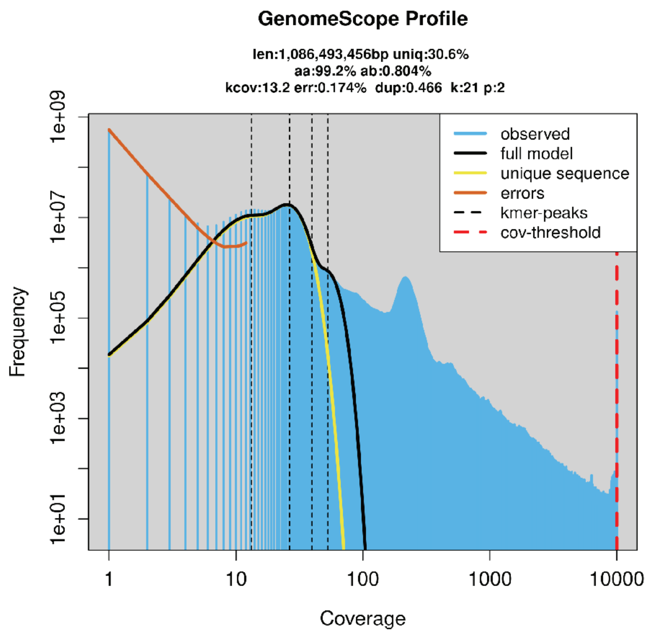

**c**

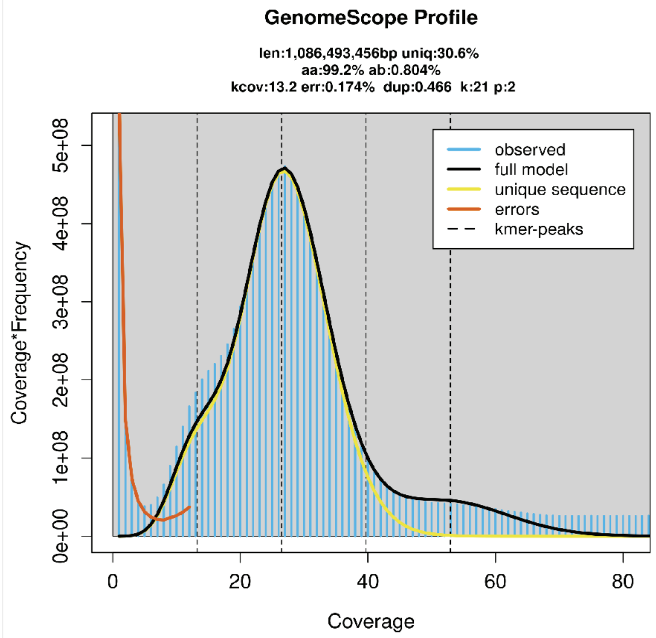

**d**

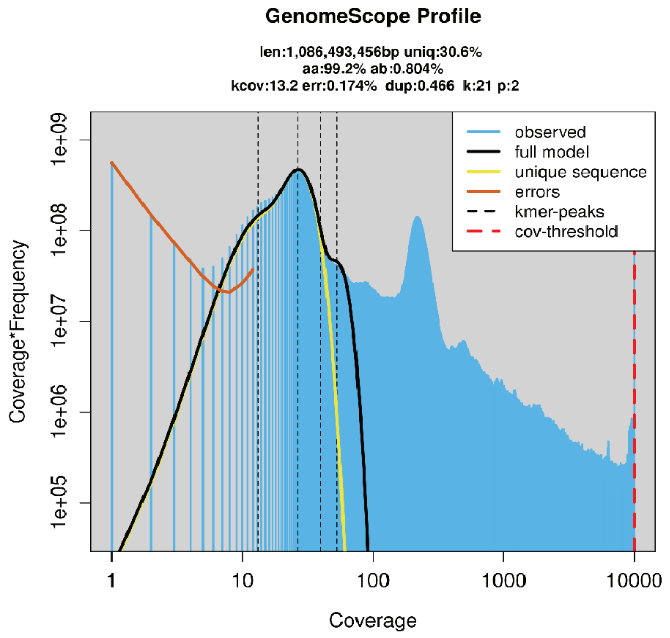

**e**

**Results**

```
GenomeScope version 2.0
input file = user_uploads/VWK4PMoK5KfpyfYrtfxl
output directory = user_data/VWK4PMoK5KfpyfYrtfxl
p = 2
k = 21

property                      min               max
Homozygous (aa)               97.838%           100%
Heterozygous (ab)             0%                2.16196%
Genome Haploid Length         1,030,435,728 bp  1,086,493,456 bp
Genome Repeat Length          715,385,879 bp    754,304,276 bp
Genome Unique Length          315,049,849 bp    332,189,180 bp
Model Fit                     33.6129%          96.0198%
Read Error Rate               0.173612%         0.173612%
```

**f**

**Model**

```
Formula: y_transform ~ x^transform_exp * length * predict2_0(r1, k, d,e
    kmercov, bias, x)

Parameters:
         Estimate Std. Error t value Pr(>|t|)
d       6.918e-02  1.133e-02   6.108 1.21e-09 ***
r1      8.043e-03  6.788e-03   1.185   0.2362
kmercov 1.324e+01  1.753e-01  75.527  < 2e-16 ***
bias    4.662e-01  1.811e-01   2.574   0.0101 *
length  3.474e+08  2.689e+07  12.922  < 2e-16 ***
---
Signif. codes:  0 '***' 0.001 '**' 0.01 '*' 0.05 '.' 0.1 ' ' 1

Residual standard error: 22280000 on 1995 degrees of freedom

Number of iterations to convergence: 7
Achieved convergence tolerance: 1.49e-08
```

### NES740 GenomeScope

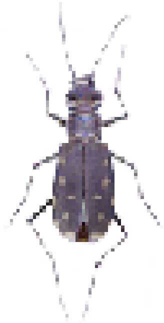

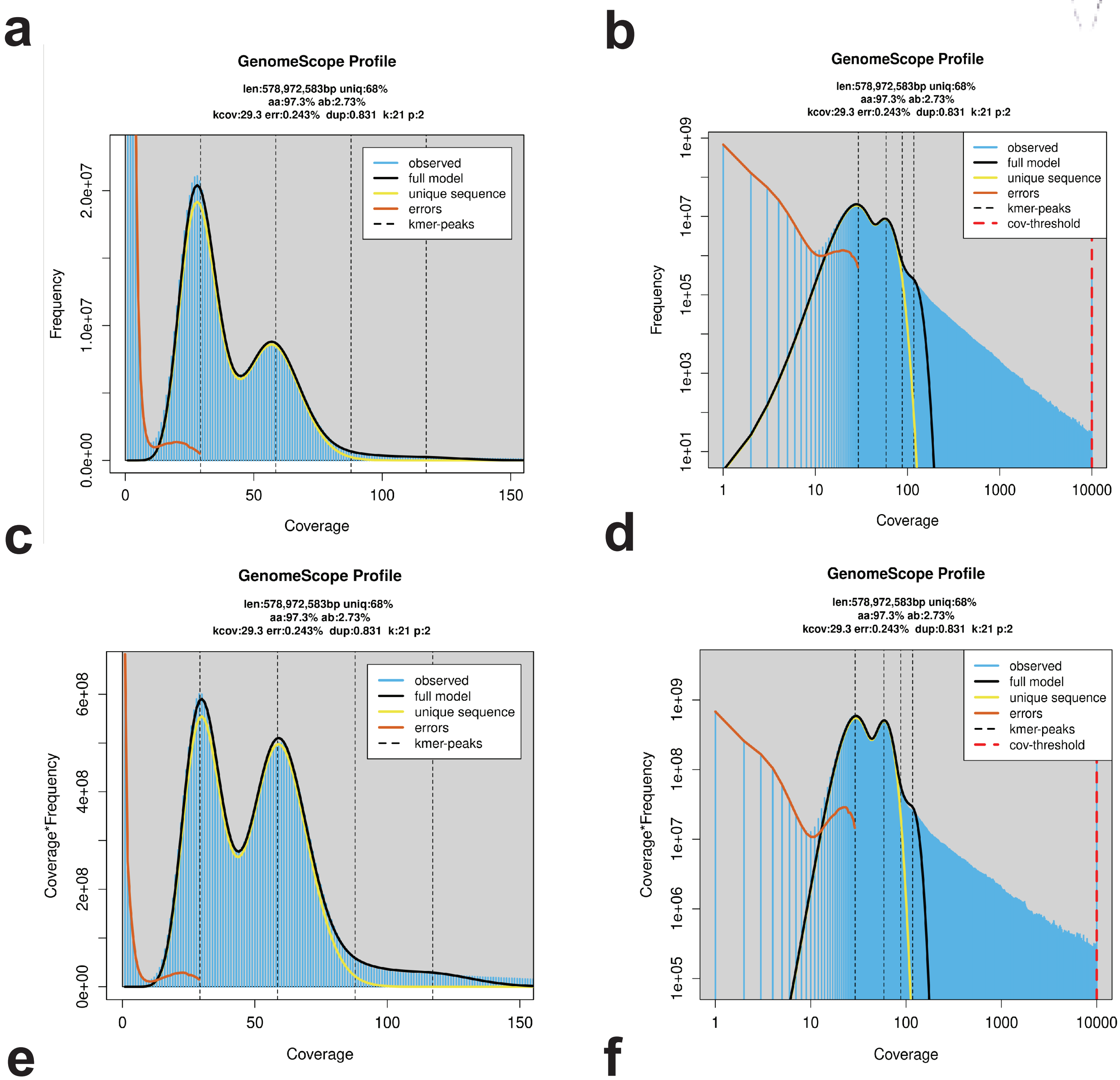

**e**

Results

```
GenomeScope version 2.0
input file = user_uploads/52DbwUA7LA1pozmaJ3cG
output directory = user_data/52DbwUA7LA1pozmaJ3cG
p = 2
k = 21

property                      min               max
Homozygous (aa)               97.2547%          97.2806%
Heterozygous (ab)             2.71942%          2.74527%
Genome Haploid Length         577,661,985 bp    578,972,583 bp
Genome Repeat Length          184,898,116 bp    185,317,613 bp
Genome Unique Length          392,763,869 bp    393,654,970 bp
Model Fit                     70.8351%          96.493%
Read Error Rate               0.242672%         0.242672%
```

**f**

Model

```
Formula: y_transform ~ x^transform_exp * length * predict2_0(r1, k, d,
    kmercov, bias, x)

Parameters:
        Estimate Std. Error t value Pr(>|t|)
d       6.056e-02  1.137e-03   53.28   <2e-16 ***
r1      2.732e-02  6.463e-05  422.75   <2e-16 ***
kmercov 2.928e+01  1.659e-02 1765.05   <2e-16 ***
bias    8.314e-01  6.919e-03  120.17   <2e-16 ***
length  4.186e+08  6.876e+05  608.74   <2e-16 ***
---
Signif. codes:  0 '***' 0.001 '**' 0.01 '*' 0.05 '.' 0.1 ' ' 1

Residual standard error: 4263000 on 1995 degrees of freedom

Number of iterations to convergence: 6
Achieved convergence tolerance: 1.49e-08
```

### NES739 GenomeScope

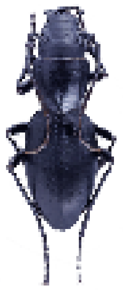

a

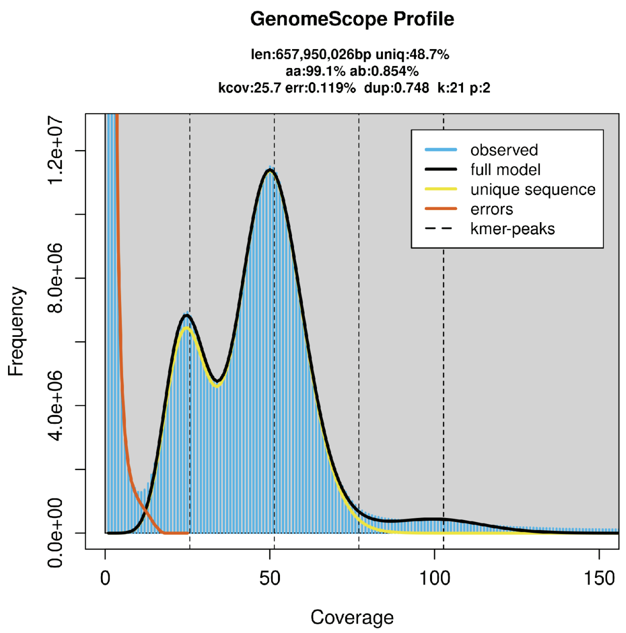

b

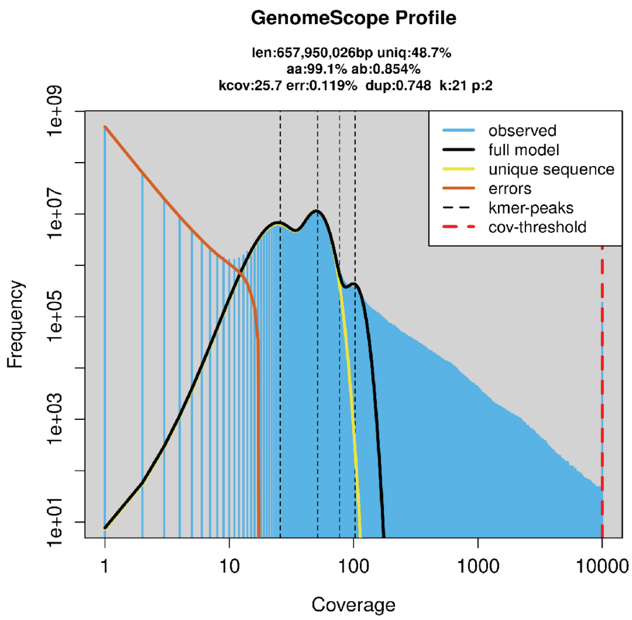

c

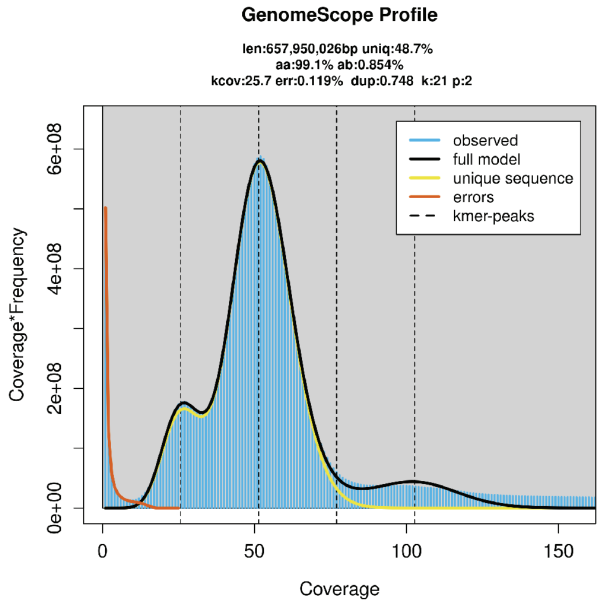

d

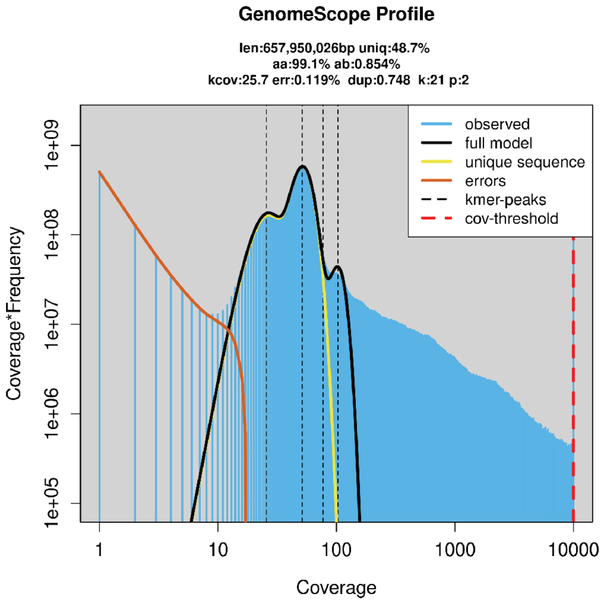

e

#### Results

```
GenomeScope version 2.0
input file = user_uploads/VkTrhxIGM6i5o9TTUjpr
output directory = user_data/VkTrhxIGM6i5o9TTUjpr
p = 2
k = 21

property                      min               max
Homozygous (aa)               99.1272%          99.1643%
Heterozygous (ab)             0.835711%         0.872777%
Genome Haploid Length         655,963,966 bp    657,950,026 bp
Genome Repeat Length          336,277,868 bp    337,296,016 bp
Genome Unique Length          319,686,098 bp    320,654,010 bp
Model Fit                     53.2302%          97.4603%
Read Error Rate               0.119245%         0.119245%
```

f

#### Model

```
Formula: y_transform ~ x^transform_exp * length * predict2_0(r1, k, d,
    kmercov, bias, x)

Parameters:
        Estimate Std. Error t value Pr(>|t|)
d       5.823e-02  1.509e-03   38.59   <2e-16 ***
r1      8.542e-03  9.266e-05   92.19   <2e-16 ***
kmercov 2.568e+01  1.941e-02 1323.14   <2e-16 ***
bias    7.475e-01  1.239e-02   60.32   <2e-16 ***
length  3.400e+08  1.038e+06  327.65   <2e-16 ***
---
Signif. codes:  0 '***' 0.001 '**' 0.01 '*' 0.05 '.' 0.1 ' ' 1

Residual standard error: 6260000 on 1995 degrees of freedom

Number of iterations to convergence: 5
Achieved convergence tolerance: 1.49e-08
```

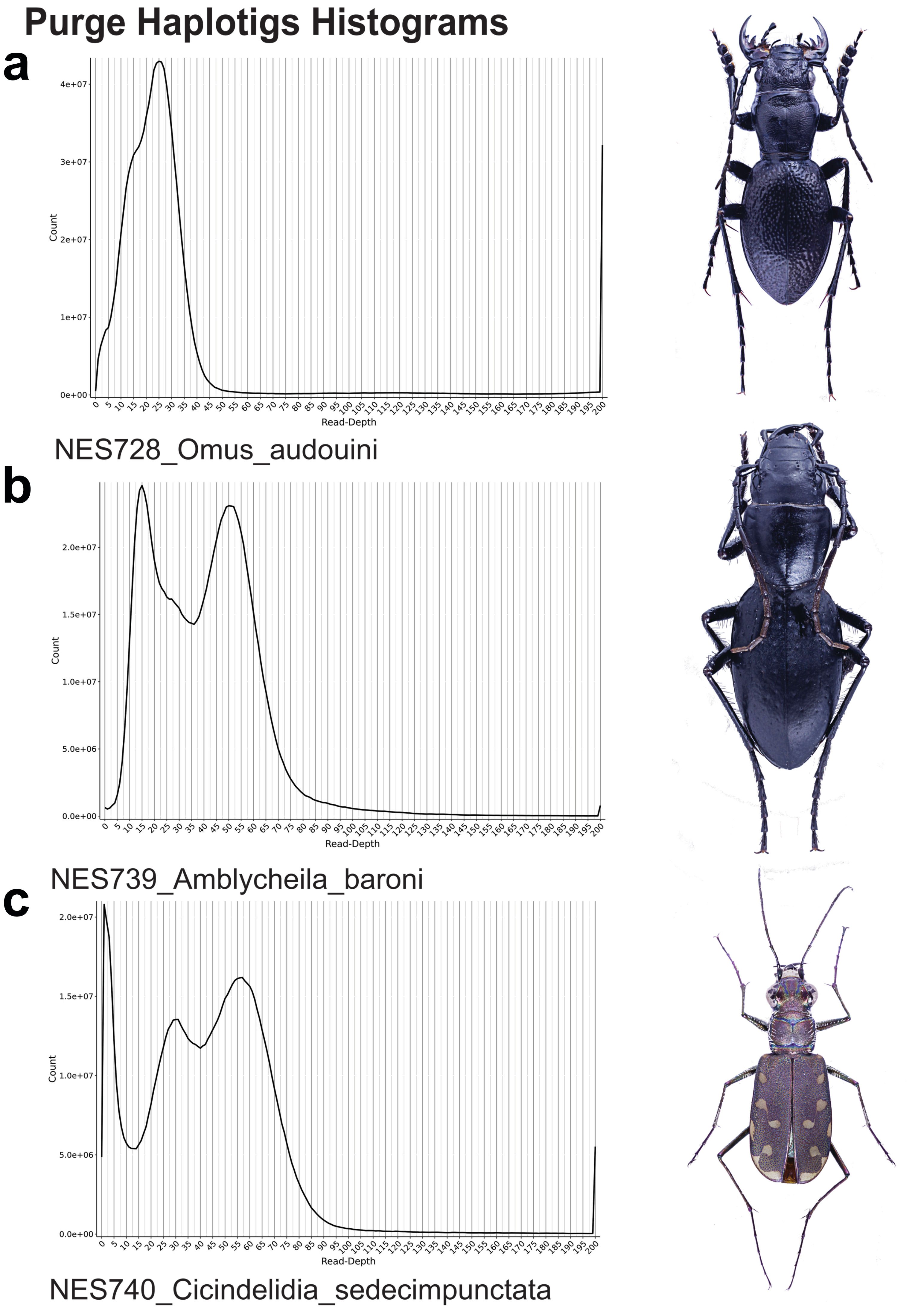
Purge Haplotigs Histograms
a
Count
Read-Depth
NES728_Omus_audouini
b
Count
Read-Depth
NES739_Amblycheila_baroni
c
Count
Read-Depth
NES740_Cicindelidia_sedecimpunctata

NES740

### Comparison of BlobTools Snailplots following different Purge haplotig purge thresholds.

a

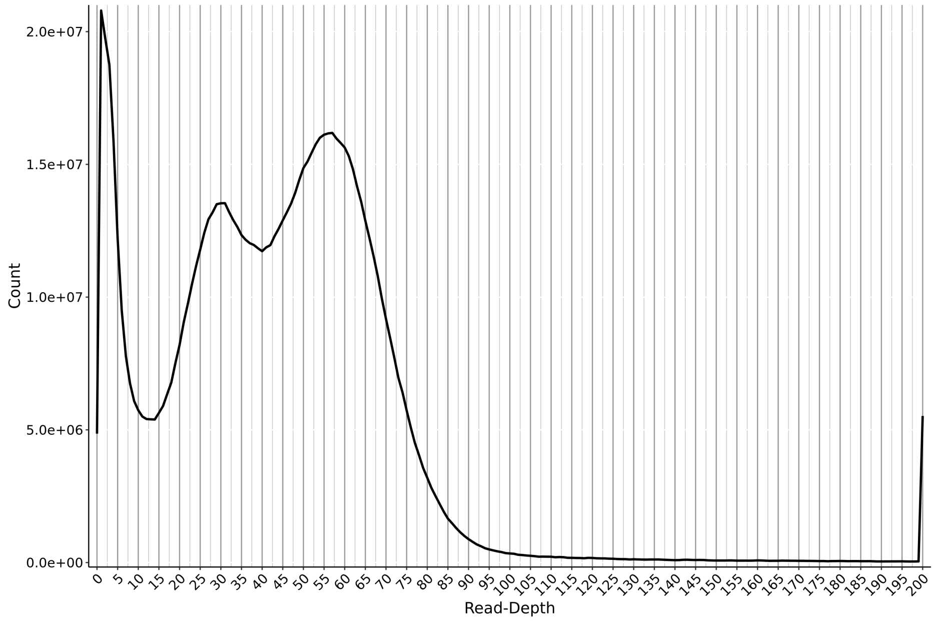

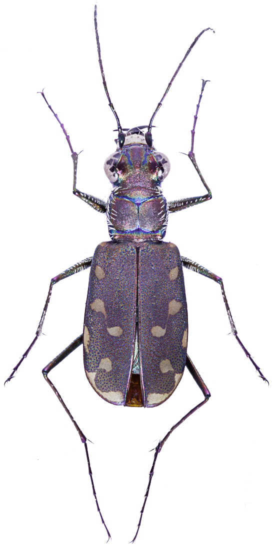

b

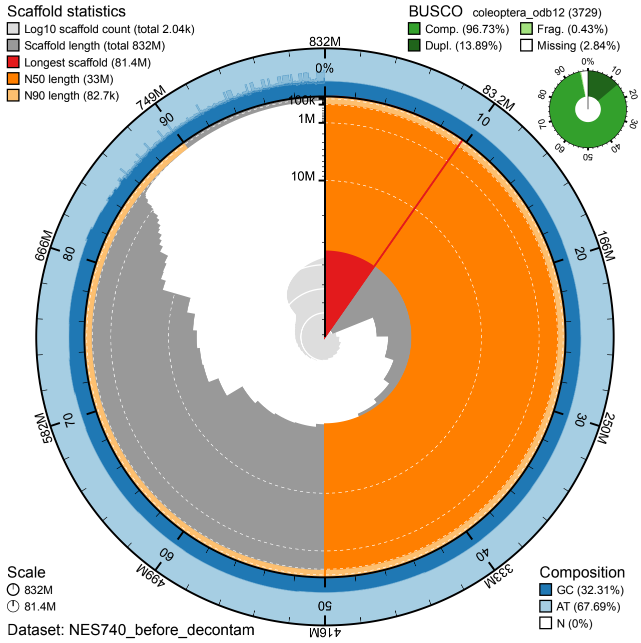

-l 10 -m 40 -h 110

c

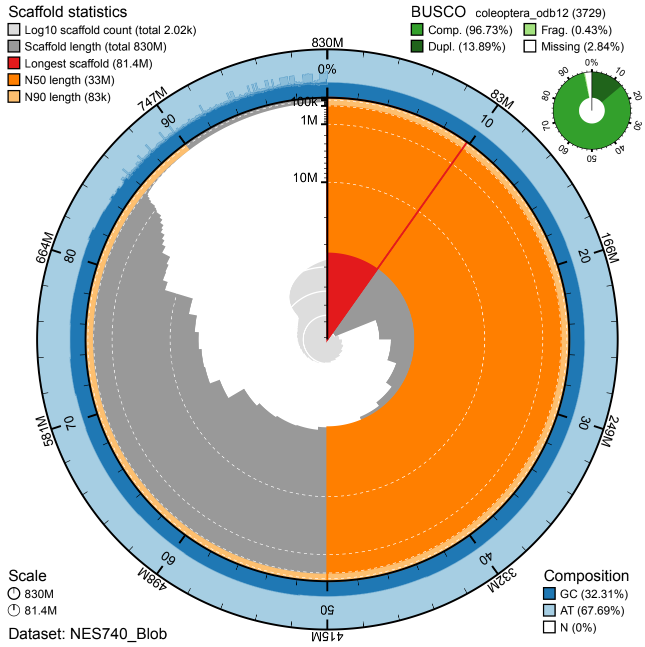

-l 10 -m 40 -h 80

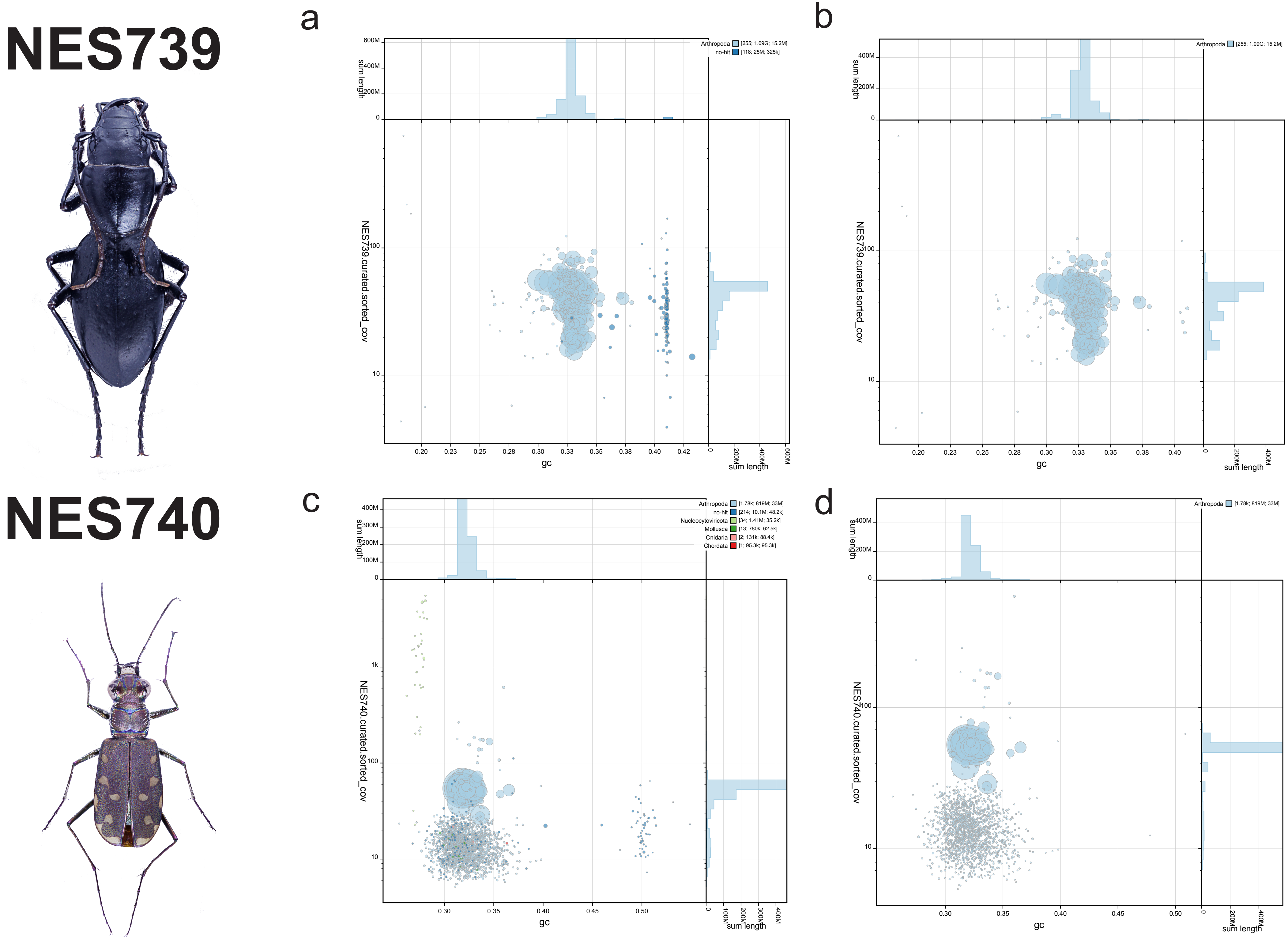
NES739
NES740
a
b
c
d
Arthropoda [255; 1.09G; 15.2M]
no-hit [118; 25M; 325k]
Arthropoda [1.78k; 819M; 33M]
no-hit [214; 10.1M; 48.2k]
Nucleocytoviricota [34; 1.41M; 35.2k]
Mollusca [13; 780k; 62.5k]
Cnidaria [2; 131k; 88.4k]
Chordata [1; 95.3k; 95.3k]
sum length
gc
NES739.curated.sorted_cov
NES740.curated.sorted_cov

### NES728

#### w/ Chordata

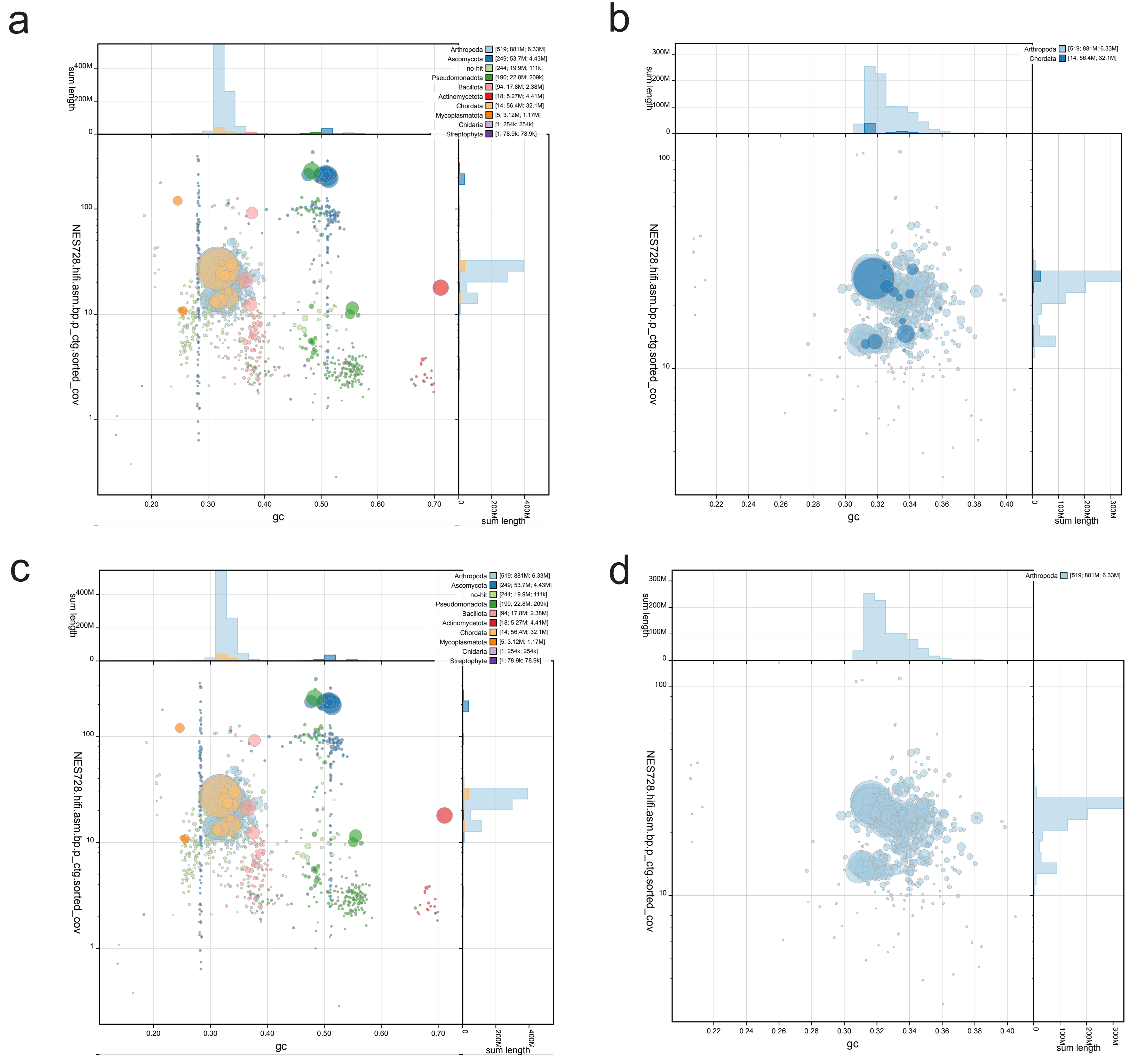

#### w/out Chordata

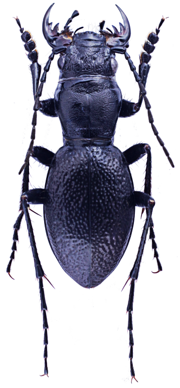

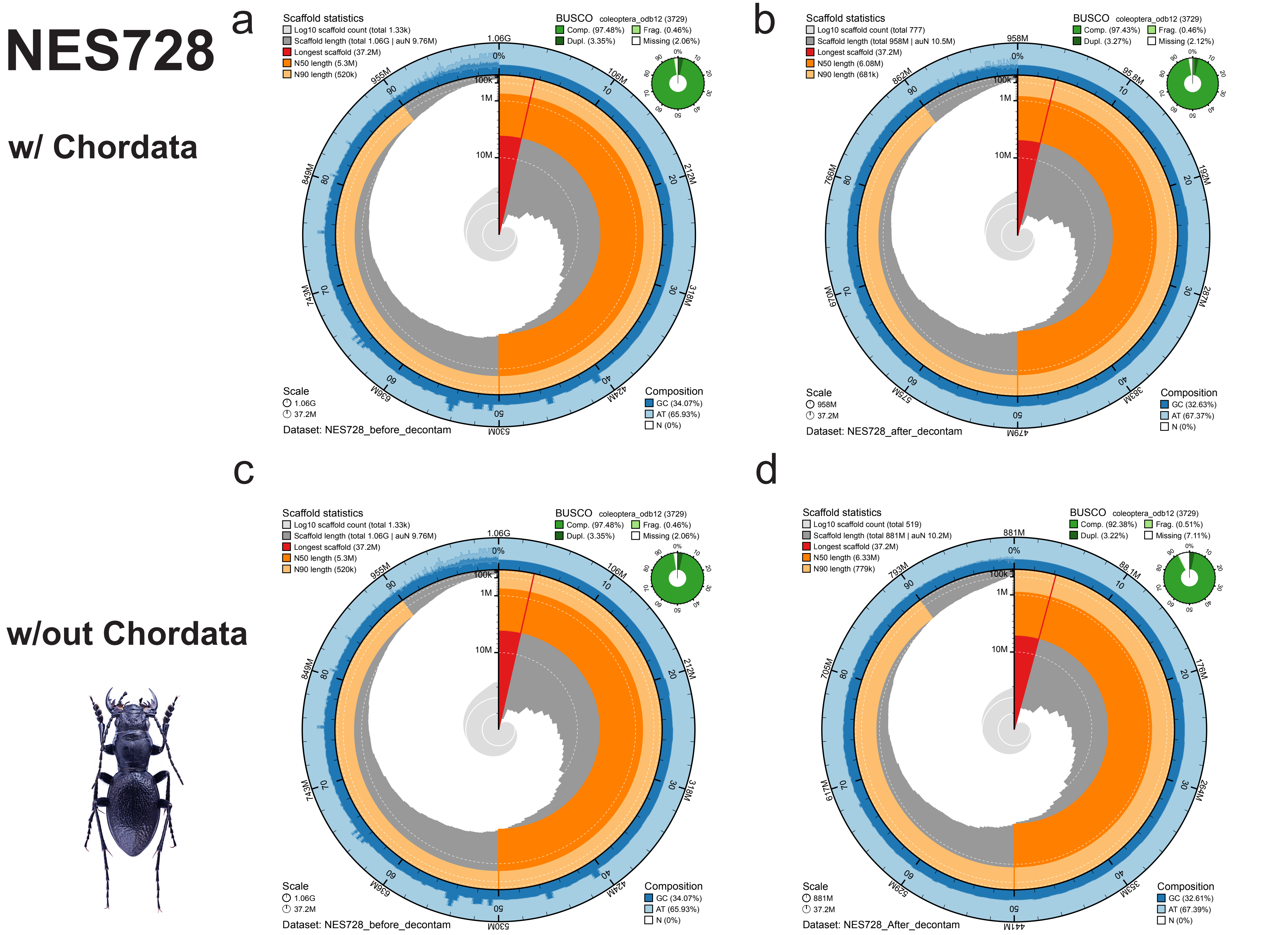
NES728
w/ Chordata
w/out Chordata
a
Scaffold statistics
Log10 scaffold count (total 1.33k)
Scaffold length (total 1.06G | auN 9.76M)
Longest scaffold (37.2M)
N50 length (5.3M)
N90 length (520k)
BUSCO coleoptera_odb12 (3729)
Comp. (97.48%) Frag. (0.46%)
Dupl. (3.35%) Missing (2.06%)
Scale
1.06G
37.2M
Composition
GC (34.07%)
AT (65.93%)
N (0%)
Dataset: NES728_before_decontam
b
Scaffold statistics
Log10 scaffold count (total 777)
Scaffold length (total 958M | auN 10.5M)
Longest scaffold (37.2M)
N50 length (6.08M)
N90 length (681k)
BUSCO coleoptera_odb12 (3729)
Comp. (97.43%) Frag. (0.46%)
Dupl. (3.27%) Missing (2.12%)
Scale
958M
37.2M
Composition
GC (32.63%)
AT (67.37%)
N (0%)
Dataset: NES728_after_decontam
c
Scaffold statistics
Log10 scaffold count (total 1.33k)
Scaffold length (total 1.06G | auN 9.76M)
Longest scaffold (37.2M)
N50 length (5.3M)
N90 length (520k)
BUSCO coleoptera_odb12 (3729)
Comp. (97.48%) Frag. (0.46%)
Dupl. (3.35%) Missing (2.06%)
Scale
1.06G
37.2M
Composition
GC (34.07%)
AT (65.93%)
N (0%)
Dataset: NES728_before_decontam
d
Scaffold statistics
Log10 scaffold count (total 519)
Scaffold length (total 881M | auN 10.2M)
Longest scaffold (37.2M)
N50 length (6.33M)
N90 length (779k)
BUSCO coleoptera_odb12 (3729)
Comp. (92.38%) Frag. (0.51%)
Dupl. (3.22%) Missing (7.11%)
Scale
881M
37.2M
Composition
GC (32.61%)
AT (67.39%)
N (0%)
Dataset: NES728_After_decontam

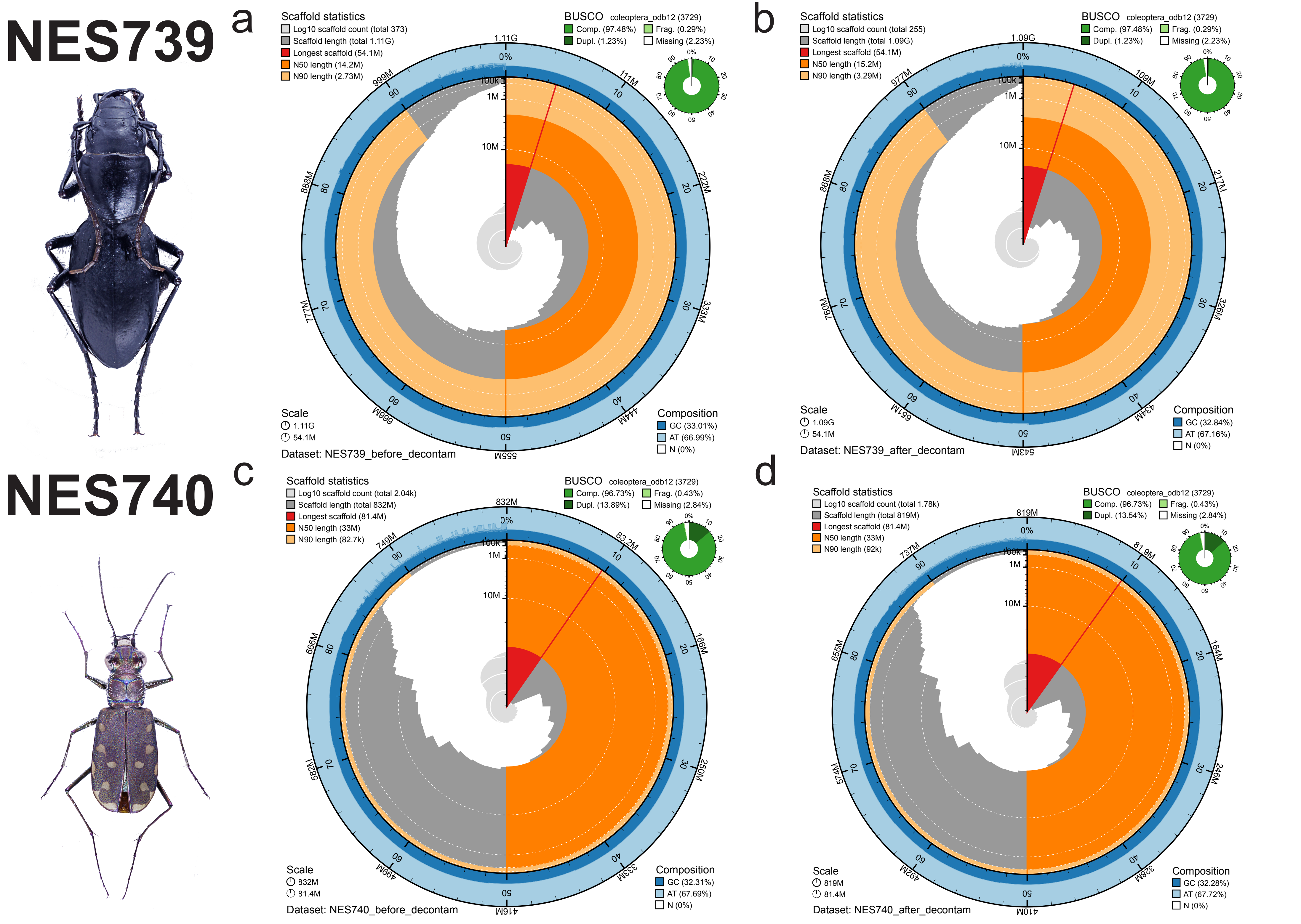
NES739
a
Scaffold statistics
Log10 scaffold count (total 373)
Scaffold length (total 1.11G)
Longest scaffold (54.1M)
N50 length (14.2M)
N90 length (2.73M)
BUSCO coleoptera_odb12 (3729)
Comp. (97.48%)
Frag. (0.29%)
Dupl. (1.23%)
Missing (2.23%)
Scale
1.11G
54.1M
Composition
GC (33.01%)
AT (66.99%)
N (0%)
Dataset: NES739_before_decontam
b
Scaffold statistics
Log10 scaffold count (total 255)
Scaffold length (total 1.09G)
Longest scaffold (54.1M)
N50 length (15.2M)
N90 length (3.29M)
BUSCO coleoptera_odb12 (3729)
Comp. (97.48%)
Frag. (0.29%)
Dupl. (1.23%)
Missing (2.23%)
Scale
1.09G
54.1M
Composition
GC (32.84%)
AT (67.16%)
N (0%)
Dataset: NES739_after_decontam
NES740
c
Scaffold statistics
Log10 scaffold count (total 2.04k)
Scaffold length (total 832M)
Longest scaffold (81.4M)
N50 length (33M)
N90 length (82.7k)
BUSCO coleoptera_odb12 (3729)
Comp. (96.73%)
Frag. (0.43%)
Dupl. (13.89%)
Missing (2.84%)
Scale
832M
81.4M
Composition
GC (32.31%)
AT (67.69%)
N (0%)
Dataset: NES740_before_decontam
d
Scaffold statistics
Log10 scaffold count (total 1.78k)
Scaffold length (total 819M)
Longest scaffold (81.4M)
N50 length (33M)
N90 length (92k)
BUSCO coleoptera_odb12 (3729)
Comp. (96.73%)
Frag. (0.43%)
Dupl. (13.54%)
Missing (2.84%)
Scale
819M
81.4M
Composition
GC (32.28%)
AT (67.72%)
N (0%)
Dataset: NES740_after_decontam

ptg000020l
ptg000027l
ptg000065l
ptg000082l
ptg000097l
ptg000104l
ptg000116l
ptg000125l
ptg000210l
ptg000272l
ptg000279l
ptg000312l
ptg000346l
ptg000523l

| | | | | | | |
|---|---|---|---|---|---|---|
| ptg000020l | 48640 | 14789 | ptg000020l | NES11_397 | 97.694 | 8630 |
| ptg000020l | 48640 | 942 | ptg000020l | NES11_397 | 97.978 | 544 |
| ptg000020l | 48640 | 486 | ptg000020l | NES11_397 | 89.817 | 383 |
| ptg000020l | 48640 | 466 | ptg000020l | NES11_397 | 88.25 | 400 |
| ptg000020l | 48640 | 462 | ptg000020l | NES11_397 | 88.119 | 404 |
| ptg000020l | 48640 | 460 | ptg000020l | NES11_397 | 88.03 | 401 |
| ptg000020l | 48640 | 455 | ptg000020l | NES11_397 | 87.841 | 403 |
| ptg000020l | 48640 | 455 | ptg000020l | NES11_397 | 87.94 | 398 |
| ptg000020l | 48640 | 453 | ptg000020l | NES11_397 | 87.719 | 399 |
| ptg000020l | 48640 | 453 | ptg000020l | NES11_397 | 87.781 | 401 |
| ptg000020l | 48640 | 449 | ptg000020l | NES11_397 | 87.909 | 397 |
| ptg000020l | 48640 | 448 | ptg000020l | NES11_397 | 87.317 | 410 |
| ptg000020l | 48640 | 444 | ptg000020l | NES11_397 | 87.657 | 397 |
| ptg000020l | 48640 | 440 | ptg000020l | NES11_397 | 87.154 | 397 |
| ptg000020l | 48640 | 440 | ptg000020l | NES11_397 | 87.406 | 397 |
| ptg000020l | 48640 | 440 | ptg000020l | NES11_397 | 87.374 | 396 |
| ptg000020l | 48640 | 438 | ptg000020l | NES11_397 | 86.978 | 407 |
| ptg000020l | 48640 | 438 | ptg000020l | NES11_397 | 86.978 | 407 |
| ptg000020l | 48640 | 435 | ptg000020l | NES11_397 | 87.831 | 378 |
| ptg000020l | 48640 | 431 | ptg000020l | NES11_397 | 86.232 | 414 |
| ptg000020l | 48640 | 431 | ptg000020l | NES11_397 | 92.739 | 303 |
| ptg000020l | 48640 | 427 | ptg000020l | NES11_397 | 92.013 | 313 |
| ptg000020l | 48640 | 427 | ptg000020l | NES11_397 | 86.553 | 409 |
| ptg000020l | 48640 | 427 | ptg000020l | NES11_397 | 86.199 | 413 |
| ptg000020l | 48640 | 425 | ptg000020l | NES11_397 | 86.52 | 408 |
| ptg000020l | 48640 | 424 | ptg000020l | NES11_397 | 87.738 | 367 |
| ptg000020l | 48640 | 422 | ptg000020l | NES11_397 | 91.909 | 309 |
| ptg000020l | 48640 | 422 | ptg000020l | NES11_397 | 86.582 | 395 |
| ptg000020l | 48640 | 422 | ptg000020l | NES11_397 | 86.432 | 398 |
| ptg000020l | 48640 | 420 | ptg000020l | NES11_397 | 92.131 | 305 |
| ptg000020l | 48640 | 418 | ptg000020l | NES11_397 | 85.714 | 406 |
| ptg000020l | 48640 | 418 | ptg000020l | NES11_397 | 91.803 | 305 |
| ptg000020l | 48640 | 416 | ptg000020l | NES11_397 | 91.586 | 309 |
| ptg000020l | 48640 | 414 | ptg000020l | NES11_397 | 91.803 | 305 |
| ptg000020l | 48640 | 414 | ptg000020l | NES11_397 | 91.531 | 307 |
| ptg000020l | 48640 | 409 | ptg000020l | NES11_397 | 85.68 | 412 |
| ptg000020l | 48640 | 409 | ptg000020l | NES11_397 | 85.645 | 411 |
| ptg000020l | 48640 | 409 | ptg000020l | NES11_397 | 91.054 | 313 |
| ptg000020l | 48640 | 407 | ptg000020l | NES11_397 | 85.442 | 419 |
| ptg000020l | 48640 | 405 | ptg000020l | NES11_397 | 85.337 | 416 |
| ptg000020l | 48640 | 405 | ptg000020l | NES11_397 | 85.372 | 417 |
| ptg000020l | 48640 | 405 | ptg000020l | NES11_397 | 90.312 | 320 |
| ptg000020l | 48640 | 403 | ptg000020l | NES11_397 | 90.735 | 313 |
| ptg000020l | 48640 | 403 | ptg000020l | NES11_397 | 90.735 | 313 |
| ptg000020l | 48640 | 401 | ptg000020l | NES11_397 | 90.536 | 317 |

| | | | | | | |
|---|---|---|---|---|---|---|
| ptg000020l | 48640 | 399 | ptg000020l | NES11_397 | 90.415 | 313 |
| ptg000020l | 48640 | 399 | ptg000020l | NES11_397 | 85.23 | 413 |
| ptg000020l | 48640 | 398 | ptg000020l | NES11_397 | 85.167 | 418 |
| ptg000020l | 48640 | 398 | ptg000020l | NES11_397 | 93.066 | 274 |
| ptg000020l | 48640 | 396 | ptg000020l | NES11_397 | 85.23 | 413 |
| ptg000020l | 48640 | 396 | ptg000020l | NES11_397 | 85.024 | 414 |
| ptg000020l | 48640 | 394 | ptg000020l | NES11_397 | 85.012 | 407 |
| ptg000020l | 48640 | 394 | ptg000020l | NES11_397 | 85.132 | 417 |
| ptg000020l | 48640 | 392 | ptg000020l | NES11_397 | 85.468 | 406 |
| ptg000020l | 48640 | 392 | ptg000020l | NES11_397 | 84.988 | 413 |
| ptg000020l | 48640 | 392 | ptg000020l | NES11_397 | 85.396 | 404 |
| ptg000020l | 48640 | 392 | ptg000020l | NES11_397 | 90.064 | 312 |
| ptg000020l | 48640 | 392 | ptg000020l | NES11_397 | 90.228 | 307 |
| ptg000020l | 48640 | 390 | ptg000020l | NES11_397 | 84.841 | 409 |
| ptg000020l | 48640 | 390 | ptg000020l | NES11_397 | 86.667 | 375 |
| ptg000020l | 48640 | 388 | ptg000020l | NES11_397 | 84.689 | 418 |
| ptg000020l | 48640 | 388 | ptg000020l | NES11_397 | 84.746 | 413 |
| ptg000020l | 48640 | 388 | ptg000020l | NES11_397 | 85.25 | 400 |
| ptg000020l | 48640 | 387 | ptg000020l | NES11_397 | 84.434 | 424 |
| ptg000020l | 48640 | 387 | ptg000020l | NES11_397 | 89.623 | 318 |
| ptg000020l | 48640 | 385 | ptg000020l | NES11_397 | 89.59 | 317 |
| ptg000020l | 48640 | 381 | ptg000020l | NES11_397 | 89.355 | 310 |
| ptg000020l | 48640 | 381 | ptg000020l | NES11_397 | 87.5 | 344 |
| ptg000020l | 48640 | 377 | ptg000020l | NES11_397 | 88.889 | 324 |
| ptg000020l | 48640 | 377 | ptg000020l | NES11_397 | 88.65 | 326 |
| ptg000020l | 48640 | 375 | ptg000020l | NES11_397 | 84.559 | 408 |
| ptg000020l | 48640 | 374 | ptg000020l | NES11_397 | 84.096 | 415 |
| ptg000020l | 48640 | 374 | ptg000020l | NES11_397 | 88.959 | 317 |
| ptg000020l | 48640 | 372 | ptg000020l | NES11_397 | 88.344 | 326 |
| ptg000020l | 48640 | 372 | ptg000020l | NES11_397 | 88.785 | 321 |
| ptg000020l | 48640 | 372 | ptg000020l | NES11_397 | 88.785 | 321 |
| ptg000020l | 48640 | 370 | ptg000020l | NES11_397 | 83.933 | 417 |
| ptg000020l | 48640 | 370 | ptg000020l | NES11_397 | 84.019 | 413 |
| ptg000020l | 48640 | 368 | ptg000020l | NES11_397 | 87.126 | 334 |
| ptg000020l | 48640 | 368 | ptg000020l | NES11_397 | 88.644 | 317 |
| ptg000020l | 48640 | 368 | ptg000020l | NES11_397 | 88.608 | 316 |
| ptg000020l | 48640 | 368 | ptg000020l | NES11_397 | 88.644 | 317 |
| ptg000020l | 48640 | 368 | ptg000020l | NES11_397 | 89.068 | 311 |
| ptg000020l | 48640 | 368 | ptg000020l | NES11_397 | 90.526 | 285 |
| ptg000020l | 48640 | 366 | ptg000020l | NES11_397 | 83.777 | 413 |
| ptg000020l | 48640 | 366 | ptg000020l | NES11_397 | 86.819 | 349 |
| ptg000020l | 48640 | 364 | ptg000020l | NES11_397 | 83.61 | 421 |
| ptg000020l | 48640 | 364 | ptg000020l | NES11_397 | 87.842 | 329 |
| ptg000020l | 48640 | 364 | ptg000020l | NES11_397 | 88.498 | 313 |
| ptg000020l | 48640 | 364 | ptg000020l | NES11_397 | 83.732 | 418 |

| | | | | | | |
|---|---|---|---|---|---|---|
| ptg000020l | 48640 | 364 | ptg000020l | NES11_397 | 83.732 | 418 |
| ptg000020l | 48640 | 364 | ptg000020l | NES11_397 | 86.389 | 360 |
| ptg000020l | 48640 | 363 | ptg000020l | NES11_397 | 83.61 | 421 |
| ptg000020l | 48640 | 363 | ptg000020l | NES11_397 | 88.328 | 317 |
| ptg000020l | 48640 | 363 | ptg000020l | NES11_397 | 84.675 | 385 |
| ptg000020l | 48640 | 361 | ptg000020l | NES11_397 | 87.963 | 324 |
| ptg000020l | 48640 | 361 | ptg000020l | NES11_397 | 84.211 | 399 |
| ptg000020l | 48640 | 361 | ptg000020l | NES11_397 | 84.04 | 401 |
| ptg000020l | 48640 | 361 | ptg000020l | NES11_397 | 88.162 | 321 |
| ptg000020l | 48640 | 359 | ptg000020l | NES11_397 | 90.288 | 278 |
| ptg000020l | 48640 | 355 | ptg000020l | NES11_397 | 83.693 | 417 |
| ptg000020l | 48640 | 355 | ptg000020l | NES11_397 | 86.217 | 341 |
| ptg000020l | 48640 | 355 | ptg000020l | NES11_397 | 87.85 | 321 |
| ptg000020l | 48640 | 355 | ptg000020l | NES11_397 | 85.278 | 360 |
| ptg000020l | 48640 | 353 | ptg000020l | NES11_397 | 83.535 | 413 |
| ptg000020l | 48640 | 353 | ptg000020l | NES11_397 | 85.146 | 377 |
| ptg000020l | 48640 | 353 | ptg000020l | NES11_397 | 87.692 | 325 |
| ptg000020l | 48640 | 351 | ptg000020l | NES11_397 | 87.538 | 329 |
| ptg000020l | 48640 | 351 | ptg000020l | NES11_397 | 87.616 | 323 |
| ptg000020l | 48640 | 351 | ptg000020l | NES11_397 | 87.898 | 314 |
| ptg000020l | 48640 | 351 | ptg000020l | NES11_397 | 90.146 | 274 |
| ptg000020l | 48640 | 350 | ptg000020l | NES11_397 | 86.866 | 335 |
| ptg000020l | 48640 | 350 | ptg000020l | NES11_397 | 86.127 | 346 |
| ptg000020l | 48640 | 348 | ptg000020l | NES11_397 | 87.385 | 325 |
| ptg000020l | 48640 | 348 | ptg000020l | NES11_397 | 87.385 | 325 |
| ptg000020l | 48640 | 346 | ptg000020l | NES11_397 | 83.292 | 407 |
| ptg000020l | 48640 | 346 | ptg000020l | NES11_397 | 86.378 | 323 |
| ptg000020l | 48640 | 346 | ptg000020l | NES11_397 | 85.755 | 351 |
| ptg000020l | 48640 | 344 | ptg000020l | NES11_397 | 86.866 | 335 |
| ptg000020l | 48640 | 344 | ptg000020l | NES11_397 | 87.421 | 318 |
| ptg000020l | 48640 | 344 | ptg000020l | NES11_397 | 87.117 | 326 |
| ptg000020l | 48640 | 344 | ptg000020l | NES11_397 | 87.227 | 321 |
| ptg000020l | 48640 | 340 | ptg000020l | NES11_397 | 82.66 | 421 |
| ptg000020l | 48640 | 340 | ptg000020l | NES11_397 | 86.93 | 329 |
| ptg000020l | 48640 | 339 | ptg000020l | NES11_397 | 86.81 | 326 |
| ptg000020l | 48640 | 339 | ptg000020l | NES11_397 | 89.781 | 274 |
| ptg000020l | 48640 | 337 | ptg000020l | NES11_397 | 86.405 | 331 |
| ptg000020l | 48640 | 337 | ptg000020l | NES11_397 | 82.506 | 423 |
| ptg000020l | 48640 | 337 | ptg000020l | NES11_397 | 82.506 | 423 |
| ptg000020l | 48640 | 337 | ptg000020l | NES11_397 | 85.673 | 342 |
| ptg000020l | 48640 | 335 | ptg000020l | NES11_397 | 86.486 | 333 |
| ptg000020l | 48640 | 335 | ptg000020l | NES11_397 | 85.455 | 330 |
| ptg000020l | 48640 | 333 | ptg000020l | NES11_397 | 81.986 | 433 |
| ptg000020l | 48640 | 333 | ptg000020l | NES11_397 | 86.503 | 326 |
| ptg000020l | 48640 | 329 | ptg000020l | NES11_397 | 82.227 | 422 |

| | | | | | | |
|---|---|---|---|---|---|---|
| ptg000020l | 48640 | 327 | ptg000020l | NES11_397 | 82.456 | 399 |
| ptg000020l | 48640 | 327 | ptg000020l | NES11_397 | 86.262 | 313 |
| ptg000020l | 48640 | 327 | ptg000020l | NES11_397 | 86.145 | 332 |
| ptg000020l | 48640 | 327 | ptg000020l | NES11_397 | 86.731 | 309 |
| ptg000020l | 48640 | 326 | ptg000020l | NES11_397 | 86.111 | 324 |
| ptg000020l | 48640 | 324 | ptg000020l | NES11_397 | 86.018 | 329 |
| ptg000020l | 48640 | 324 | ptg000020l | NES11_397 | 86.068 | 323 |
| ptg000020l | 48640 | 324 | ptg000020l | NES11_397 | 88.321 | 274 |
| ptg000020l | 48640 | 322 | ptg000020l | NES11_397 | 82.057 | 418 |
| ptg000020l | 48640 | 322 | ptg000020l | NES11_397 | 86.018 | 329 |
| ptg000020l | 48640 | 320 | ptg000020l | NES11_397 | 81.56 | 423 |
| ptg000020l | 48640 | 320 | ptg000020l | NES11_397 | 85.897 | 312 |
| ptg000020l | 48640 | 320 | ptg000020l | NES11_397 | 85.759 | 316 |
| ptg000020l | 48640 | 313 | ptg000020l | NES11_397 | 81.542 | 428 |
| ptg000020l | 48640 | 309 | ptg000020l | NES11_397 | 85.231 | 325 |
| ptg000020l | 48640 | 309 | ptg000020l | NES11_397 | 89.243 | 251 |
| ptg000020l | 48640 | 307 | ptg000020l | NES11_397 | 84.94 | 332 |
| ptg000020l | 48640 | 307 | ptg000020l | NES11_397 | 81.818 | 418 |
| ptg000020l | 48640 | 298 | ptg000020l | NES11_397 | 81.01 | 416 |
| ptg000020l | 48640 | 296 | ptg000020l | NES11_397 | 80.841 | 428 |
| ptg000020l | 48640 | 296 | ptg000020l | NES11_397 | 84.545 | 330 |
| ptg000020l | 48640 | 294 | ptg000020l | NES11_397 | 91.031 | 223 |
| ptg000020l | 48640 | 292 | ptg000020l | NES11_397 | 84.179 | 335 |
| ptg000020l | 48640 | 292 | ptg000020l | NES11_397 | 88.048 | 251 |
| ptg000020l | 48640 | 291 | ptg000020l | NES11_397 | 81.281 | 406 |
| ptg000020l | 48640 | 289 | ptg000020l | NES11_397 | 88.477 | 243 |
| ptg000020l | 48640 | 287 | ptg000020l | NES11_397 | 87.649 | 251 |
| ptg000020l | 48640 | 285 | ptg000020l | NES11_397 | 91.549 | 213 |
| ptg000020l | 48640 | 285 | ptg000020l | NES11_397 | 84.295 | 312 |
| ptg000020l | 48640 | 285 | ptg000020l | NES11_397 | 87.6 | 250 |
| ptg000020l | 48640 | 285 | ptg000020l | NES11_397 | 88.115 | 244 |
| ptg000020l | 48640 | 283 | ptg000020l | NES11_397 | 87.5 | 256 |
| ptg000020l | 48640 | 283 | ptg000020l | NES11_397 | 88.115 | 244 |
| ptg000020l | 48640 | 281 | ptg000020l | NES11_397 | 84.127 | 315 |
| ptg000020l | 48640 | 281 | ptg000020l | NES11_397 | 87.251 | 251 |
| ptg000020l | 48640 | 281 | ptg000020l | NES11_397 | 87.5 | 248 |
| ptg000020l | 48640 | 276 | ptg000020l | NES11_397 | 81.471 | 367 |
| ptg000020l | 48640 | 276 | ptg000020l | NES11_397 | 86.853 | 251 |
| ptg000020l | 48640 | 276 | ptg000020l | NES11_397 | 86.853 | 251 |
| ptg000020l | 48640 | 274 | ptg000020l | NES11_397 | 82.029 | 345 |
| ptg000020l | 48640 | 272 | ptg000020l | NES11_397 | 83.183 | 333 |
| ptg000020l | 48640 | 270 | ptg000020l | NES11_397 | 82.934 | 334 |
| ptg000020l | 48640 | 270 | ptg000020l | NES11_397 | 86.454 | 251 |
| ptg000020l | 48640 | 270 | ptg000020l | NES11_397 | 86.454 | 251 |
| ptg000020l | 48640 | 268 | ptg000020l | NES11_397 | 84.266 | 286 |

| index | identifiers | gc | length | bestsumorder_phylum |
|---|---|---|---|---|
| 0 | ptg000020l | 0.3177 | 32051766 | Arthropoda |
| 1 | ptg000027l | 0.3374 | 6081416 | Arthropoda |
| 2 | ptg000065l | 0.3401 | 1589396 | Arthropoda |
| 3 | ptg000082l | 0.3257 | 3475717 | Arthropoda |
| 4 | ptg000097l | 0.3336 | 944574 | Arthropoda |
| 5 | ptg000104l | 0.3418 | 2408723 | Arthropoda |
| 6 | ptg000116l | 0.3355 | 795940 | Arthropoda |
| 7 | ptg000125l | 0.3184 | 4227880 | Arthropoda |
| 8 | ptg000210l | 0.3243 | 610285 | Arthropoda |
| 9 | ptg000272l | 0.3475 | 338057 | Arthropoda |
| 10 | ptg000279l | 0.3301 | 1755750 | Arthropoda |
| 11 | ptg000312l | 0.3366 | 232356 | Arthropoda |
| 12 | ptg000346l | 0.3126 | 1733138 | Arthropoda |
| 13 | ptg000523l | 0.3589 | 191569 | Arthropoda |

### Quast Results: NES728

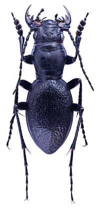

a

#### Chordata Filtered

Report

| | NES728_hifi_filtered_no_Chordata |
|---|---|
| # contigs (>= 0 bp) | 519 |
| # contigs (>= 1000 bp) | 519 |
| # contigs (>= 5000 bp) | 519 |
| # contigs (>= 10000 bp) | 519 |
| # contigs (>= 25000 bp) | 516 |
| # contigs (>= 50000 bp) | 481 |
| Total length (>= 0 bp) | 881420873 |
| Total length (>= 1000 bp) | 881420873 |
| Total length (>= 5000 bp) | 881420873 |
| Total length (>= 10000 bp) | 881420873 |
| Total length (>= 25000 bp) | 881361108 |
| Total length (>= 50000 bp) | 880089875 |
| # contigs | 519 |
| Largest contig | 37178479 |
| Total length | 881420873 |
| GC (%) | 32.61 |
| N50 | 6329733 |
| N90 | 778888 |
| auN | 10154757.5 |
| L50 | 32 |
| L90 | 209 |
| # N's per 100 kbp | 0.00 |

All statistics are based on contigs of size >= 500 bp, unless otherwise noted (e.g., "# contigs (>= 0 bp)" and "Total length (>= 0 bp)" include all contigs).

b

#### With Chordata

Report

| | NES728_hifi_filtered_with_Chordata |
|---|---|
| # contigs (>= 0 bp) | 533 |
| # contigs (>= 1000 bp) | 533 |
| # contigs (>= 5000 bp) | 533 |
| # contigs (>= 10000 bp) | 533 |
| # contigs (>= 25000 bp) | 530 |
| # contigs (>= 50000 bp) | 495 |
| Total length (>= 0 bp) | 937857440 |
| Total length (>= 1000 bp) | 937857440 |
| Total length (>= 5000 bp) | 937857440 |
| Total length (>= 10000 bp) | 937857440 |
| Total length (>= 25000 bp) | 937797675 |
| Total length (>= 50000 bp) | 936526442 |
| # contigs | 533 |
| Largest contig | 37178479 |
| Total length | 937857440 |
| GC (%) | 32.60 |
| N50 | 6759456 |
| N90 | 795940 |
| auN | 10728056.7 |
| L50 | 32 |
| L90 | 214 |
| # N's per 100 kbp | 0.00 |

All statistics are based on contigs of size >= 500 bp, unless otherwise noted (e.g., "# contigs (>= 0 bp)" and "Total length (>= 0 bp)" include all contigs).

c

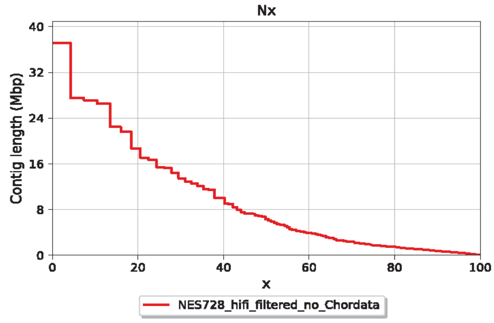

d

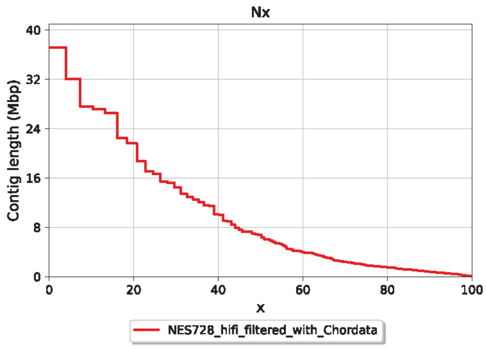

e

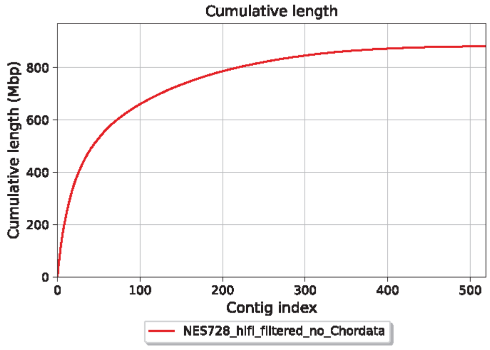

f

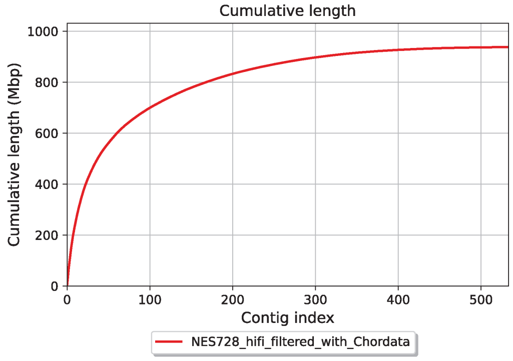

g

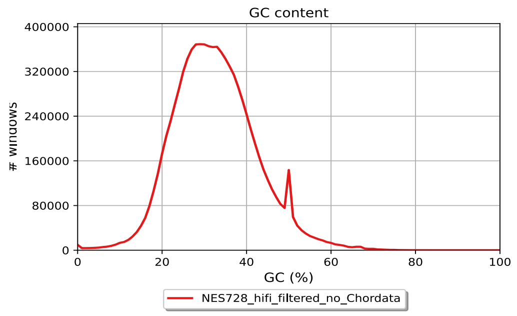

h

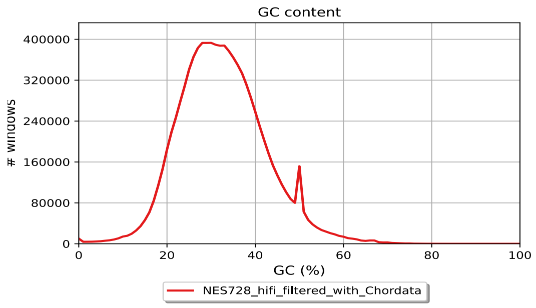

i

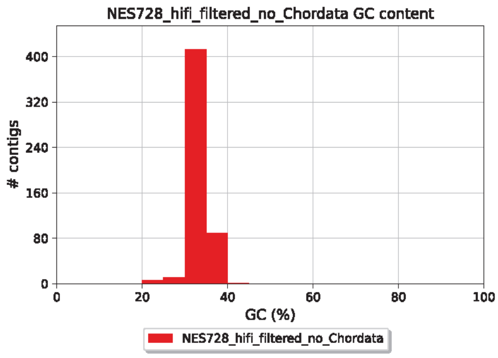

j

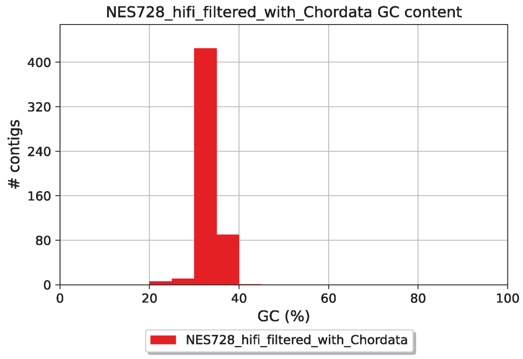

### Quast Results

a

NES739 Report

| | NES739_hifi_filtered_no_hit |
|---|---|
| # contigs (>= 0 bp) | 255 |
| # contigs (>= 1000 bp) | 255 |
| # contigs (>= 5000 bp) | 255 |
| # contigs (>= 10000 bp) | 255 |
| # contigs (>= 25000 bp) | 252 |
| # contigs (>= 50000 bp) | 239 |
| Total length (>= 0 bp) | 1085518503 |
| Total length (>= 1000 bp) | 1085518503 |
| Total length (>= 5000 bp) | 1085518503 |
| Total length (>= 10000 bp) | 1085518503 |
| Total length (>= 25000 bp) | 1085461019 |
| Total length (>= 50000 bp) | 1084978652 |
| # contigs | 255 |
| Largest contig | 54143589 |
| Total length | 1085518503 |
| GC (%) | 32.84 |
| N50 | 15194716 |
| N90 | 3287415 |
| auN | 18294794.7 |
| L50 | 22 |
| L90 | 79 |
| # N's per 100 kbp | 0.00 |

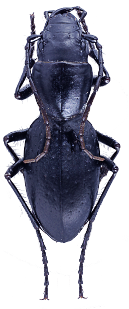

All statistics are based on contigs of size >= 500 bp, unless otherwise noted (e.g., "# contigs (>= 0 bp)" and "Total length (>= 0 bp)" include all contigs).

b

NES740 Report

| | NES740_hifi_filtered_no_contam |
|---|---|
| # contigs (>= 0 bp) | 1780 |
| # contigs (>= 1000 bp) | 1780 |
| # contigs (>= 5000 bp) | 1780 |
| # contigs (>= 10000 bp) | 1780 |
| # contigs (>= 25000 bp) | 1772 |
| # contigs (>= 50000 bp) | 1065 |
| Total length (>= 0 bp) | 819365743 |
| Total length (>= 1000 bp) | 819365743 |
| Total length (>= 5000 bp) | 819365743 |
| Total length (>= 10000 bp) | 819365743 |
| Total length (>= 25000 bp) | 819182718 |
| Total length (>= 50000 bp) | 791225240 |
| # contigs | 1780 |
| Largest contig | 81424639 |
| Total length | 819365743 |
| GC (%) | 32.28 |
| N50 | 33048812 |
| N90 | 92016 |
| auN | 35743849.3 |
| L50 | 9 |
| L90 | 260 |
| # N's per 100 kbp | 0.00 |

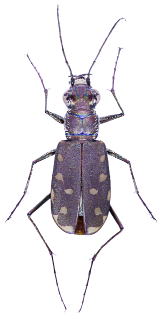

All statistics are based on contigs of size >= 500 bp, unless otherwise noted (e.g., "# contigs (>= 0 bp)" and "Total length (>= 0 bp)" include all contigs).

c

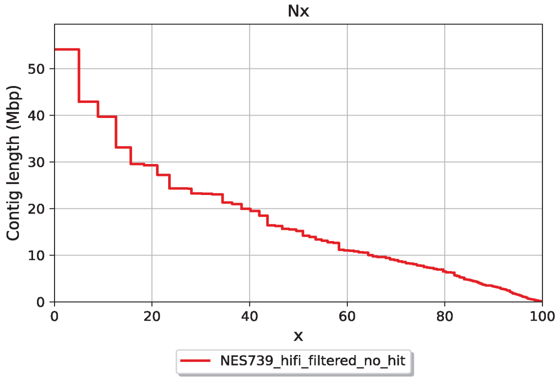

d

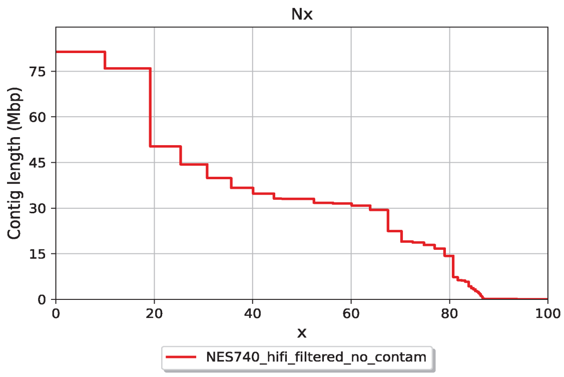

e

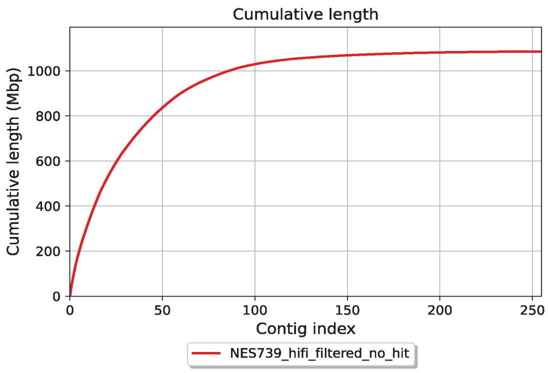

f

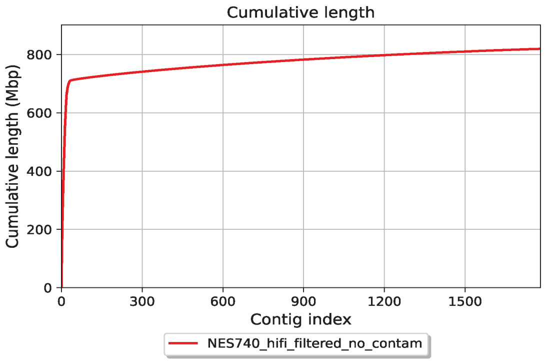

g

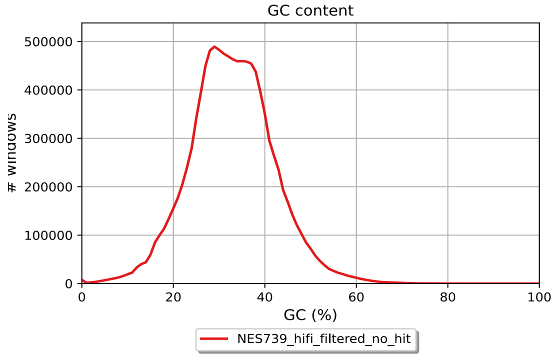

h

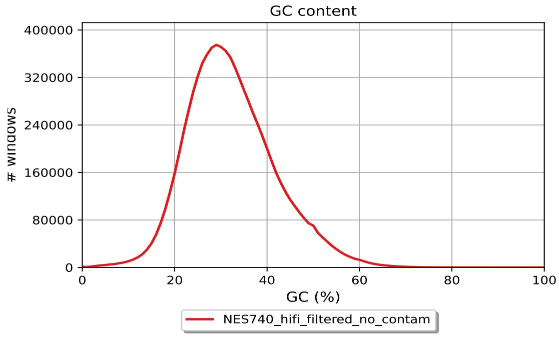

i

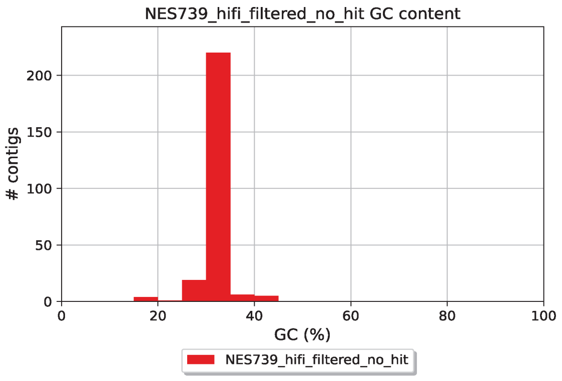

j

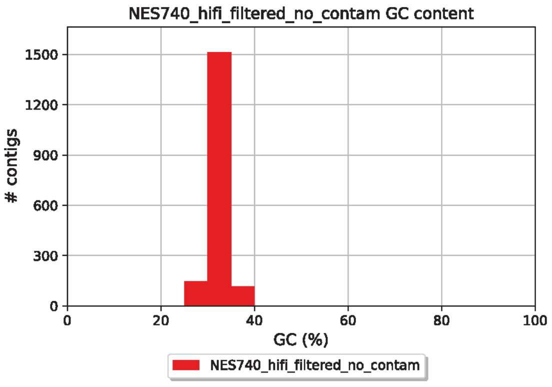
